# Learning the structural diversity in random protein sequence space

**DOI:** 10.64898/2026.04.30.722084

**Authors:** Filip Buchel, Tereza Neuwirthova, Theodora Tureckiova, Gustavo Fuertes, Ales Benda, Dalibor Panek, Matus Fricek, Mohammed AlQuraishi, Klara Hlouchova

**Affiliations:** Department of Cell Biology, Faculty of Science, Charles University, Prague, Czech Republic; Department of Biochemistry, Faculty of Science, Charles University, Prague, Czech Republic; Institute of Biotechnology of the Czech Academy of Sciences, Vestec, Czech Republic; Department of Systems Biology, Columbia University, New York, NY, USA; Institute of Organic Chemistry and Biochemistry of the Czech Academy of Sciences, Prague, Czech Republic; Imaging Methods Core Facility at BIOCEV, Vestec, Czech Republic

## Abstract

The universe of possible protein sequences is astronomically large, yet our understanding of sequence-structure relationships is derived almost entirely from the infinitesimal fraction used by life. Determining which physical properties are already accessible to unevolved polypeptides, and which require subsequent evolutionary refinement, is central both to understanding protein emergence and to expanding protein design beyond biological sequence space. Here, we explore this largely uncharted space using a high-throughput FRET-based selection of a library containing millions of random protein sequences, complemented by fluorescence imaging, biophysical characterization and sequence-based modelling. We find that random sequence space contains multiple physical phenotypes that cannot be captured by a simple distinction between soluble and aggregated proteins. Individual sequences can produce substantial FRET in both the cytosol and visible assemblies, while assemblies differ in packing density and cellular responses. Repeated selection for stronger FRET reduces population solubility indicating that proximity-based selection alone does not necessarily favor biology-like protein states. These experimentally defined phenotypes are predictable from sequence.

Our results reveal resolvable and learnable variation within unevolved protein sequence space, providing a route to investigate how far random sequences lie from soluble, stable protein solutions and to learn from regions of the protein universe not previously sampled by evolution.

## Introduction

Protein sequences occupy an astronomically large combinatorial space, and learning to navigate this space is a central challenge in protein design. Most contemporary advances in structure prediction and *de novo* protein design rely on naturally occurring sequences produced by evolutionary exploration of sequence space ^1–4^. However, available sequence annotations and experimentally solved structures are unevenly distributed, with strong biases toward a limited set of organisms and protein families ^5^. As a result, large regions of sequence space - used by biology, evolutionary dormant or entirely unexplored - remain sparsely sampled, potentially limiting the generality of current design principles.

Random sampling of protein sequence space provides a complementary, orthogonal strategy for exploring structure-function relationships. Selections from random sequence libraries have yielded proteins with diverse functions, including ligand binding and enzymatic activity ^6–8^. Random polypeptide libraries have been used to train predictors of aggregation propensity, underscoring the potential of probing global sequence-structure relationships beyond evolutionary constraints ^9^. Finally, Tretyachenko and colleagues employed random libraries with defined amino acid compositions to separate populations with distinct solubility and structural content using proteolytic resistance assays ^10^. While these studies established that random sequence space contains measurable variation in solubility, structural propensity and aggregation behavior, the relationships among these properties remain difficult to resolve at scale.

Here, we systematically explore the conformational and assembly properties of a large random protein sequence library using an *in vivo* FRET-based proximity biosensor ^11,12^. After establishing how the assay reports on fluorophore proximity and intermolecular packing using control proteins, we apply it to high-throughput screening of more than one million random sequences. We combine spatially resolved fluorescence measurements with reporter-free expression to examine how intracellular localization, molecular proximity and solubility vary among the recovered sequences and populations. This allows us to interpret the selected phenotypes beyond their FRET signals and to test how repeated selection changes the properties of the recovered populations. We further examine whether the experimentally defined FRET classes can be predicted from sequences. These analyses explore the physical behaviours accessible to unevolved polypeptides and extend the experimental basis for understanding protein emergence and guiding design beyond natural sequence datasets.

## Results

### FRET measurements and fluorescence imaging report on protein conformation and assembly

FRET efficiency depends on fluorophore proximity and orientation, and FRET-based sensors have previously been used to monitor conformational changes^11,13–15^ and aggregation mechanisms in natural proteins ^16,17^. Here, we adapted a genetically encoded FRET-based sensor to investigate structural diversity in a library of random protein sequences. High-throughput detection of FRET is attainable by flow cytometry and we first resolved how apparent FRET measured by cytometry maps to conformational and assembly properties of tested proteins using time-resolved spectroscopy and fluorescence imaging. We used control proteins with previously characterized properties to build a reference for interpreting these complementary readouts.

The control proteins were expressed in *E. coli* as terminal fusions to a genetically encoded FRET pair, such that FRET efficiency (*E*_FRET_) reports on the proximity of the fluorophores, defined by structural and oligomeric properties of the insert . We tested the system using a panel of controls, including glycine-serine repeat (GS) linkers of varying length, length-matched globular *E. coli* proteins, and previously characterized^18^ random proteins. The system was characterized on multiple levels: *in vitro* assessment of *E*_FRET_ in cell lysates and on purified proteins using time-resolved fluorescence spectroscopy; *in vivo* measurements by fluorescence lifetime imaging microscopy (FLIM) to assess *E*_FRET_ dependence on localization, and flow cytometry measurement of apparent FRET to simulate the high-throughput regime. As expected, *E*_FRET_ decreased with increasing GS linker length^19^ and displayed uniform intracellular distributions. For globular *E. coli* proteins, *E*_FRET_ correlated with protein length only partially and distinct efficiencies were observed for proteins of similar size (for example, IscA, BolA, and CspA; Figure S1a-c), although the scale of the inconsistency can be attributed to their structural details (e.g. precise termini position, oligomeric state).

To establish how the sensor responds to random proteins with different structural and aggregation properties, we examined three previously characterized sequences ^18^. The disordered control 665 was soluble and showed low FRET efficiency and diffuse intracellular fluorescence (Figure 1a,b). By contrast, two controls with higher structural content, 1259 and 1856, both formed inclusion bodies (IBs) but produced substantially different FRET signals (Figure 1a,b). Protein 1259 exhibited low FRET efficiency, while 1856 showed high FRET efficiency.

**Figure 1.**
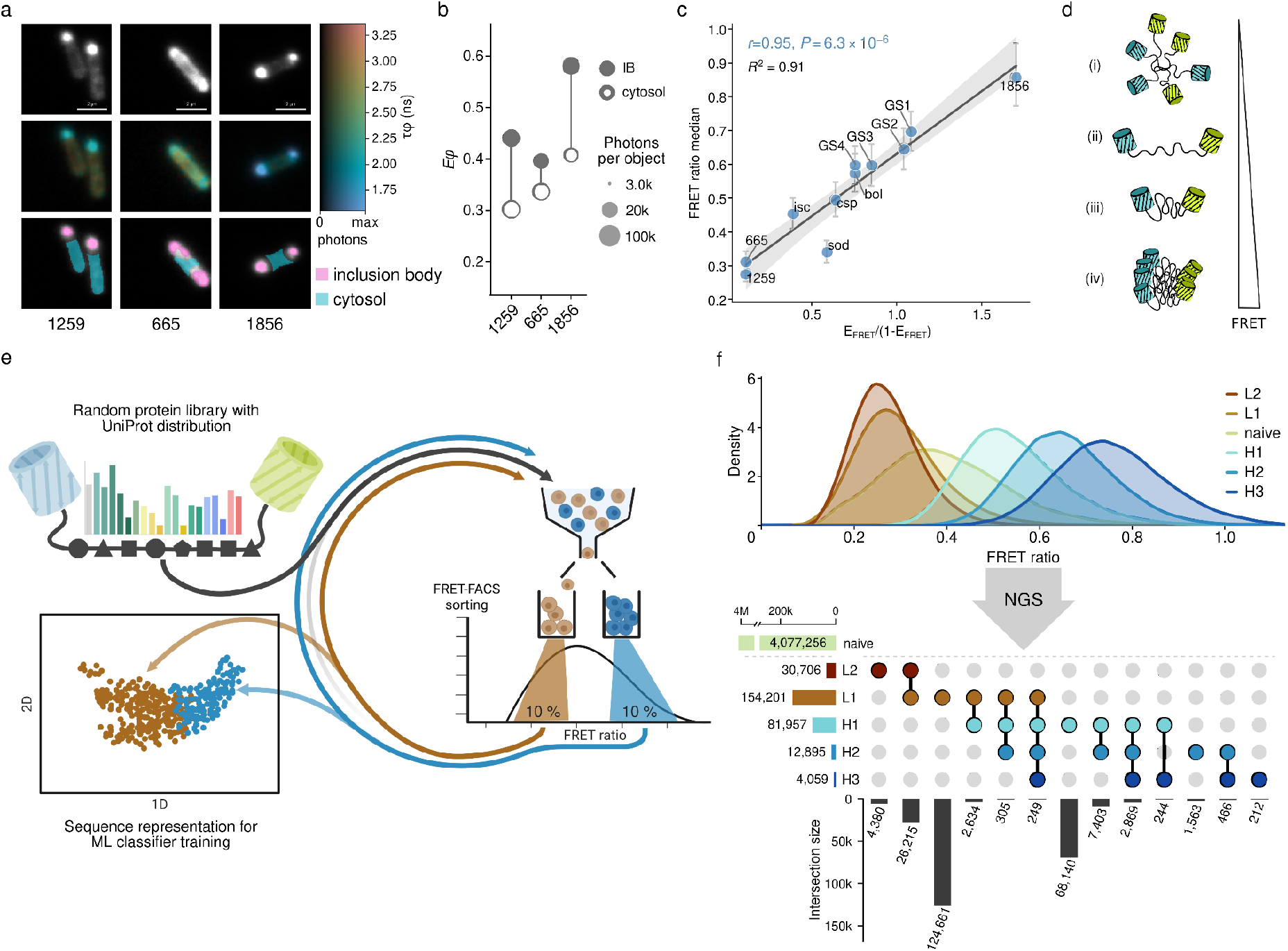
Decomposition of random protein library into structurally distinct subpopulations by iterative FRET-based screening. **a, b, c**, Characterization of the FRET-based reporter with random protein controls (disordered protein 665 and structurally enriched proteins 1259 and 1856), various length glycine-serine linkers and globular *E. coli* proteins (bolA, cspA, iscA, sodA). The controls were expressed in fusion with a FRET pair in *E. coli* and the FRET efficiency was assessed by flow cytometry, fluorescence lifetime imaging and time-resolved spectroscopy. Panel (**a**) shows FLIM images of the random protein controls. Top row — intensity images of both fluorescent channels (mVenus and mTurquoise2); middle row — intensity modulated fluorescence lifetime maps with hue encoding the phase lifetime (τ_φ_) and the brightness encoding the photon count per pixel; bottom row—cell segmentation strategy (ROI masks overlaid with intensity images). Fluorescence lifetimes of segmented regions were quantified by phasor analysis and used to calculate the FRET efficiencies (E_φ_) from E_φ_ = 1 - τ_φ_ / τ_D_ (**b**). Panel (**c**) shows correlation of cytometry-derived parameter FRET ratio and FRET efficiencies measured in cell-lysates by time-resolved spectroscopy (Whiskers show interquartile range of the FRET ratio distribution; line is least-squares linear fit; grey band is bootstrapped 95% confidence interval (1000 resamples); *r* is Pearson correlation coefficient with two-sided *P*; *R*^2^ is coefficient of determination). **d**, FRET-based proximity biosensor reports intramolecular distance of the donor and the acceptor (extended vs. compact; (ii) and (iii)), and intermolecular distances in oligomers (loosely vs. tightly packed; (i) and (iv)). Depicted scenarios map in order (i)-(iv) to increasing FRET ratio. **e**, Scheme of the screening pipeline. Random protein library with UniProt amino acid composition fused to FRET pair is expressed in *E. coli* and sorted based on FRET ratio signal by FACS in multiple rounds. Composition of the sorted subpopulations is tracked by sequencing and the data are used to extract sequence features and to train a binary classifier. **f**, Flow cytometry profiles and sequence abundance in sorted subpopulations. KDE plot shows FRET ratio of the subpopulations recorded after the sorting campaign (L1 and L2 - two rounds of low FRET sorting; H1, H2 and H3 - three rounds of high FRET sorting; naive - pre-selection library). Proportions and intersections of the sorted populations extracted from next-generation sequencing (NGS) are shown in the upset plot. The sequences were filtered based on read counts from NGS (*Methods*).

To investigate whether the contrasting FRET signals reflected differences in molecular packing, we combined FLIM with time-resolved fluorescence anisotropy measurements of the acceptor. Anisotropy measurements can detect homoFRET, i.e. energy transfer between closely spaced, identical fluorophores, through rapid loss of fluorescence polarization ^20^. Because each fusion contains only a single acceptor, homoFRET between acceptors reports specifically on proximity between different fusion molecules.

Protein 1856 showed a pronounced decrease in acceptor anisotropy within IBs, accompanied by high heteroFRET efficiency. By contrast, IBs formed by 1259 retained higher acceptor anisotropy and low FRET efficiency (Figure S1f). These observations support more ordered packing of the fluorescent reporters in 1856 assemblies than in those formed by 1259, and suggest that intermolecular energy transfer contributes to the high FRET efficiency of 1856. Thus, the contrasting signals reflect differences in the organization of the assemblies, rather than IB formation alone.

Complementary characterization of the purified controls supported the predominantly monomeric state of 665, while full-length 1259 and 1856 were recovered in high-molecular-weight fractions (Figure S1e). However, the FRET efficiency of purified 1259 construct increased substantially, indicating that its FRET signal depends on the sample complexity (Figure S1c,d).

Finally, we compared the flow-cytometric FRET ratio with FRET efficiencies determined from time-resolved measurements on cell lysates. The flow-cytometric readout closely tracked the differences among the control constructs (Figure 1c), supporting its use for high-throughput screening. Together, these experiments established a reference for interpreting FRET signals in random proteins (Figure 1d). Low FRET was observed both in a soluble, dispersed control and in an IB-forming control, whereas the high FRET of protein 1856 was associated with closer intermolecular packing of the fluorescent reporters. Thus, the FRET ratio provided a scalable screening readout, while fluorescence imaging and anisotropy measurements helped distinguish the molecular states underlying these signals.

### High-throughput FRET screening reveals diverse phenotypes within a random sequence library

To sample the sequence space of small proteins, we generated a library with 109 randomized positions and a target amino acid composition reflecting average UniProt frequencies. The library sequences were designed using the CoLiDe algorithm, previously applied to bulk studies of large random libraries with defined compositional constraints ^10,21^. To enable efficient library construction at this scale, we developed a Golden Gate Assembly-based strategy that builds the library from two oligonucleotides while preserving full diversity at the ligation junction. This approach allows cost-effective, PCR-free library generation (*Methods*). The resulting library (termed 20F109) comprised approximately four million unique variants .

We applied the FRET-based biosensor to enrich low- and high-FRET subpopulations through successive rounds of fluorescence-activated cell sorting (FACS). In the first round, cells were collected from approximately the lowest and highest 10% of the FRET-ratio distribution (Figure 1e). These populations were subjected to further sorting in the same direction, giving two low-FRET rounds (L1 and L2) and three high-FRET rounds (H1–H3). Progressive shifts in the recovered FRET-ratio distributions demonstrated enrichment of contrasting FRET phenotypes (Figure 1f). We then used next-generation sequencing (NGS) to track sequence abundance across the sorted subpopulations. Reads were mapped to a reference database generated from the naive library, allowing us to follow enrichment across successive rounds. Approximately one quarter of the naive library members were recovered in at least one sorting round, with sequence diversity progressively decreasing in later rounds (Figure 1f).

To characterize the phenotypes enriched by sorting, we selected high-abundance variants from each round for individual analysis (Figure 2a-d). Flow cytometry showed that these candidates spanned nearly the full range of FRET ratios observed in the naive library (Figure 2d). FLIM measurements then revealed how these signals related to intracellular localization, with cytosolic and inclusion body regions analyzed separately. Low-FRET candidates showed two different patterns; some displayed diffuse intracellular fluorescence, resembling the disordered control 665, whereas others formed localized assemblies with relatively low FRET efficiencies, resembling control protein 1259 (Figure 2a–c). High-FRET candidates displayed inclusion bodies with higher FRET efficiencies, resembling the behaviour of control protein 1856. However, their FRET signals were not confined to these assemblies. Several candidates (mainly variants from H1 and H2) also retained substantial FRET in cytosolic regions outside the inclusion bodies. Where both regions were analyzed, FRET efficiencies were generally higher in inclusion bodies than in the cytosol, with the difference varying among constructs (Figure 2c). Thus, individual characterization of candidates revealed variation in both intracellular localization and the FRET properties of cytosolic and assembly material.

**Figure 2.**
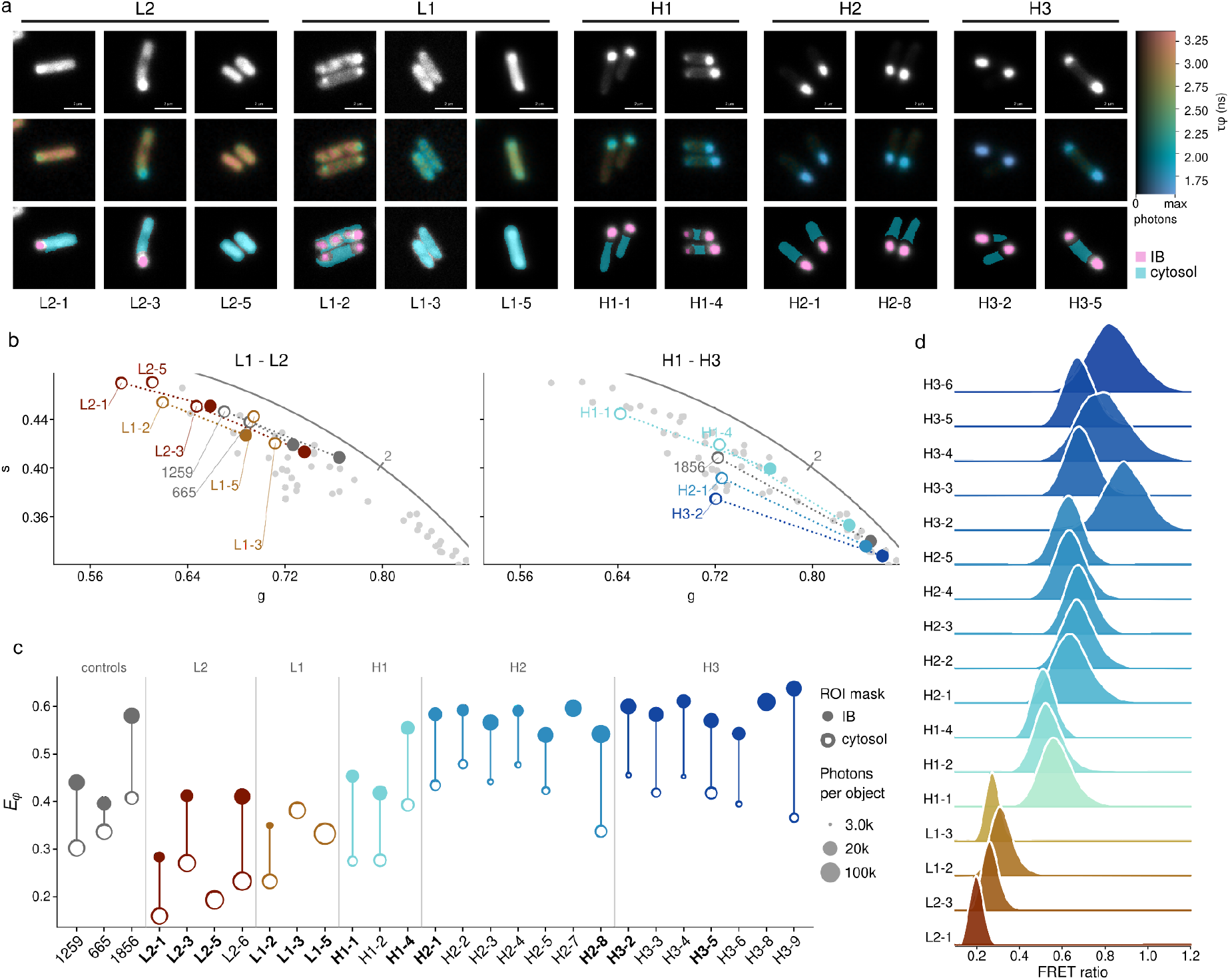
Variants identified in FRET-FACS screening recapitulate phenotypes defined by the proximity biosensor. **a**, FRET-FLIM images show localization of the fluorescent signal and per pixel donor lifetime (lifetime decrease indicates energy migration via FRET). Representative phenotypes from each sorting round are shown. Top row — intensity images of both fluorescent channels (mVenus and mTurquoise2); middle row — intensity modulated fluorescence lifetime maps with hue encoding the phase lifetime (τ_φ_) and the brightness encoding the photon count per pixel; bottom row — cell segmentation strategy (ROI masks overlaid with intensity images). Intensity images are overlaid with cytosol (cyan) and inclusion body (IB, pink) ROI masks. **b**, Summed phasors per mask type of variants shown in (**a**) (colored by selection round) and control proteins (grey). Hollow circles correspond to cytosol and full circles to IB masks. **c**, Dumbbell plot of FRET efficiencies (E_φ_) calculated from *E_φ_ = 1 - τ_φ_ / τ_D_* . Hollow circles depict cytosol and full circles IB masks, hue encodes number of photons per object. **d**, FRET ratio of selected variants measured by flow cytometry (KDE plots of FRET ratio histograms).

We next asked whether these differences in intracellular organization were accompanied by differences in cellular stress response. We monitored activation of the chaperone-associated *ibpA* promoter using a GFP reporter ^22^. Reporter activity was elevated in low-FRET candidates with localized fluorescence and in control protein 1259, but also varied among high-FRET candidates. Candidates from H3 generally showed lower responses than those from H1 and H2, while the natural protein controls and the disordered control 665 showed low reporter activity (Figure S2a). These results indicate that localized assemblies differed not only in their FRET properties but also in the cellular response detected by the *ibpA* reporter.

Together, FRET-based sorting and individual-candidate characterization revealed variation in intracellular distribution, assembly-associated FRET signals and *ibpA* activation within the random sequence library. The complementary measurements provided a reference for interpreting the sorted populations beyond their FRET ratios alone.

### FRET-selected populations differ in sequence properties and solubility

To identify sequence features associated with the observed FRET phenotypes, we extracted a set of primary sequence descriptors, including amino acid composition, hydrophobicity, isoelectric point and sequence complexity. A prominent trend was observed in hydrophobicity, which gradually decreased towards the high-FRET population H3, accompanied by enrichment of charged residues (arginine, glutamate and aspartate) (Figure 3a and Figure S3c).

**Figure 3.**
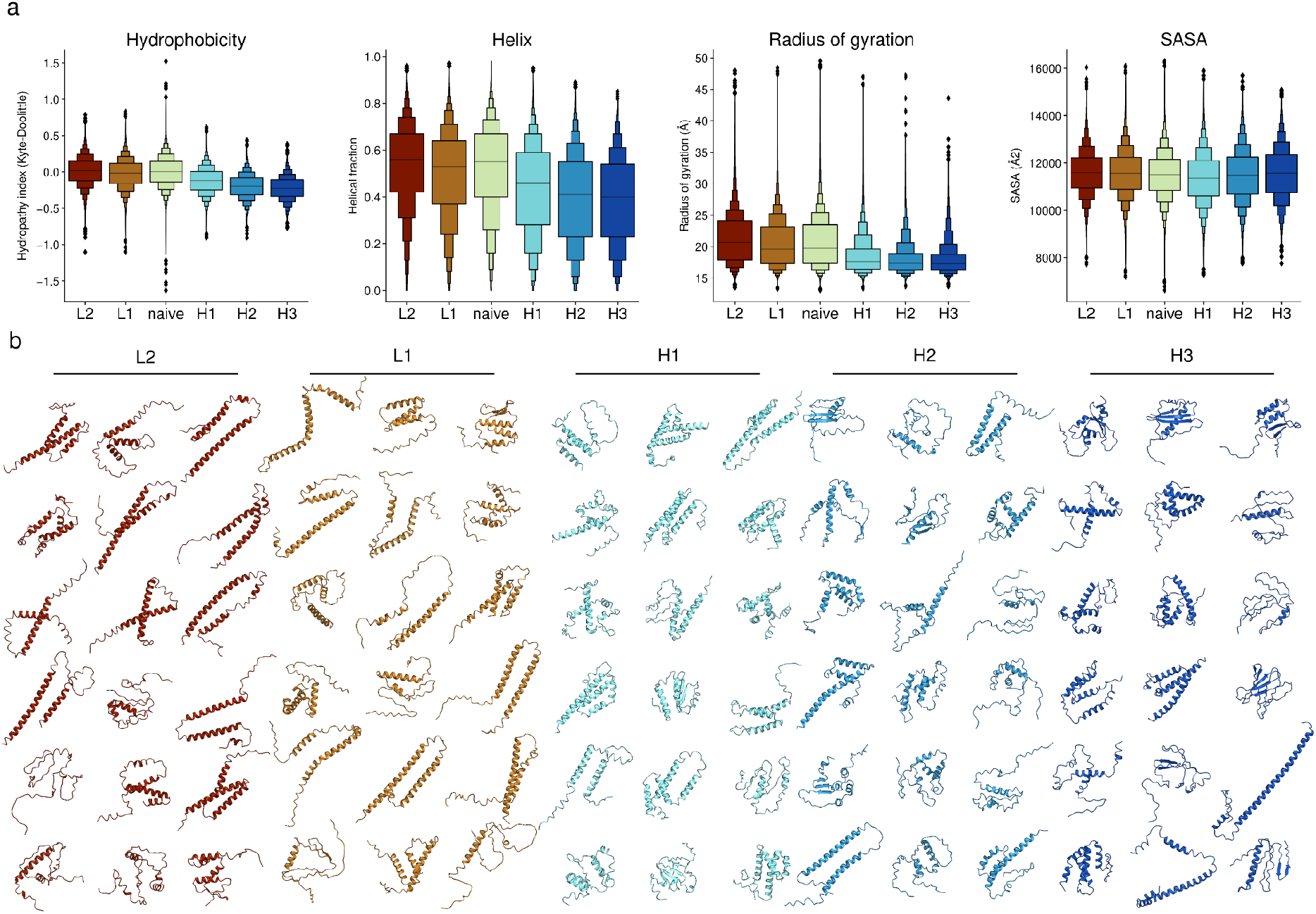
FRET-defined structural and assembly states are mirrored in sequence and structural features. **a**, Properties of subpopulations isolated in the FRET-FACS screening. Average hydrophobicity was based on the Kyte-Doolittle scale, and structural features were extracted from AlphaFold2 prediction by DSSP4 (helix - proportion of summed α-helix (H), 3_10_-helix (G), π-helix (I); Radius of gyration (Å); ASA - accessible surface area). The populations were filtered based on NGS read counts (see Methods). **b**, Predicted structures of the 18 most abundant variants from each subpopulation.

To assess the solubility of sequences enriched by FRET-based sorting, we expressed the sorted subpopulations without the FRET reporters in an *E. coli*-based reconstituted cell-free system. Coding sequences were excised from the pETMF vector and expressed with a C-terminal FLAG tag, and total and soluble protein fractions were evaluated by dot-blot assay (Methods). Relative to the naive library, all FRET-sorted subpopulations exhibited increased expression yields, although the size difference between the naive and sorted libraries might be a contributing factor. Triton X-100 generally improved expression yields of the sorted subpopulations. Low-FRET subpopulations exhibited higher solubility, whereas the fraction of soluble proteins decreased progressively across successive high-FRET selection rounds (Figure S2b).

We also extracted structural features from AlphaFold2 and ESMfold models, including secondary structure content, radius of gyration and solvent-accessible surface area (Figure 3a and Figure S2d). Prediction confidence was low, with pLDDT values around 40. While these comparisons thus describe low-confidence model-derived trends (rather than experimentally established conformations), the overall trends find good agreement between the AlphaFold2 and ESMfold models (Figure S3a). Models of low-FRET variants (L1 and L2) were predominantly extended and helical, with larger radii of gyration and greater solvent exposure, whereas models of high-FRET variants (H1–H3) were more compact and had increased β-sheet content (Figure 3a and Figure S2d).

To visualize the predicted structural variability, we selected 18 representative sequences from each sorted subpopulation and examined their AlphaFold2 structural models, prioritizing highly abundant variants (Figure 3b). Comparison of these models with structures in the PDB and AlphaFold Database using Foldseek^23^ did not identify strong matches (Figure S3b). However, the low confidence of the input models limits the interpretation of these structural comparisons.

Across successive rounds of high-FRET selection, the radius of gyration and solvent-accessible surface area of the predicted structures remained largely unchanged. However, hydrophobicity and predicted helical content decreased progressively from the first to the third high-FRET round (H1 to H3; Figure 3a), a trend that coincided with reduced solubility, as observed in cell-free expression assays (Figure S2b). Together, these results suggest that repeated selection for high FRET progressively enriches aggregation-prone variants, without a corresponding increase in predicted compactness. Earlier high-FRET populations, by contrast, retain a larger soluble fraction when expressed without the fluorescent reporters.

### Sequence-based models predict FRET-defined phenotypes

A central goal of protein design is not only to characterize individual regions of sequence space, but to learn sequence features that enable navigation across this space. We therefore asked whether the FRET-defined phenotypes leave identifiable signatures in the primary sequence that can be captured by modern protein language models.

We first compared representative sequences from the low- and high-FRET 20F109 subpopulations with length-matched natural proteins using two-dimensional projections of embeddings generated by the pretrained ESM-2 model ^24^. Despite the library being designed to reflect average natural amino acid frequencies, the selected random sequences formed clusters separated from the sampled natural proteins in the UMAP projection (Figure S3c).

We next examined whether the low- and high-FRET subpopulations could also be distinguished within the random sequence dataset. Their ESM-2 embeddings were projected into two dimensions, using the experimental FRET labels only for the projection. Despite strong overlap, sequences from the two subpopulations showed tendency to separate in 2D embedding space (Figure 3a). This suggested that language model representations might capture the sequence features associated with the experimentally defined FRET phenotypes. This behaviour was observed with the dimension reduction of the generated embeddings as much as to only two dimensions, which is a significant loss of captured information. The clearest separation tendency between the two subpopulations in 2D space was observed with mean embeddings projected with UMAP (Supplementary figures 4a-f).

Encouraged by these results, we trained supervised binary classifiers to predict FRET-defined structural classes directly from sequence. Before training, sequences were filtered to remove duplicates between subpopulations and encoded using a sequence-representation strategy (Methods). We evaluated six distinct classifier architectures, ranging from random forests to neural networks, using different encoding strategies (Figure 3 b,c). Given the overlap observed between different sorting rounds in the 2D embedding space (Supplementary figure 4g and 4h), sequences from all rounds were pooled for training (Methods). To mitigate overfitting and prevent data leakage, we employed an MMseqs2-guided diversity strategy for our train/test split, as proposed by Bushuiev & Bushuiev ^25,26^. Specifically, we clustered all 94,000 sequence variants at 50% sequence identity and coverage, ranked clusters by their mean identity to sequences outside the cluster using an all-vs-all sequence comparison (Methods), and withheld the 20% most distant clusters in their entirety (7,699 of 38,497 clusters; 11,410 variants, 12.1% of the dataset) as an independent test set, ensuring that no variant from a held-out cluster appeared in training. Because entire clusters were held out, train and test sets share no closely related variants within or across classes.

All models evaluated on these withheld clusters showed strong classification performance, with high ROC-AUC and average precision values overall, confirming robust discrimination between classes (Figure 3d, Supplementary figure 5b-e). The highest ROC-AUC was obtained with the classifier using per-token ESM-2 embeddings (Model 2, ROC-AUC = 0.91), closely followed by the neural network using one-hot encoding (Model 3, ROC-AUC = 0.90). Threshold-dependent analyses (Methods, Supplementary figure 5a) further indicated that the best operating points clustered around 0.3-0.5, highlighting that performance differences between models are most relevant when selecting deployment thresholds and calibration requirements rather than in raw accuracy alone.

Reliability-diagram analysis (Supplementary figure 5c) shows that all used models were generally close to the diagonal, however closer inspection of the calibration metrics (Log loss, Brier score and ECE (Expected Calibration Error), Supplementary figure 5d) confirms the previously observed trend on ROC curves that Model 2 and Model 3 seem to be the best performing ones. These models have the lowest ECE and Brier score which suggest better calibration than the other models. The Log loss confirms the trends for Model 2, but for the Model 3 the Log loss score is the worst from all six compared models.

To determine how accurately global physicochemical properties alone could predict the FRET-defined phenotypes, we trained a logistic-regression model on descriptors derived from each sequence (Figure 3b). The features included hydrophobicity (GRAVY), net charge at pH 7, isoelectric point (pI), ratios of amino acids grouped by their physicochemical properties, and sequence complexity (Shannon entropy) (Methods). These descriptors provided substantial predictive information (ROC-AUC = 0.77, Figure 3d) but their performance remained below that of the strongest sequence-based models. To quantify the contribution of individual feature groups, we trained separate logistic regression models on each group alone and evaluated performance on the modeling set (Figure 3e). The strongest single-group predictors were the fraction of acidic residues (ROC-AUC = 0.71) and hydrophobicity features (ROC-AUC = 0.71), whereas models based on basic, sulphur-containing, or amide residues performed close to chance (ROC-AUC ≈ 0.51). Isoelectric point and net charge alone reached moderate predictive power (ROC-AUC = 0.64 and 0.65, respectively), and Shannon entropy contributed little on its own (ROC-AUC = 0.53). To assess feature importance within the full multivariate model, we computed permutation importance as the mean decrease in ROC-AUC after random shuffling of each feature (30 repeats; Figure 3f). Isoelectric point and GRAVY were the dominant contributors (ΔROC-AUC = 0.16 and 0.13, respectively), with smaller but detectable effects for the fractions of basic and aromatic residues (ΔROC-AUC ≈ 0.015), whereas net charge, sequence complexity, and most amino-acid-group fractions had negligible impact once other features were included. These analyses identified hydrophobicity and charge-related properties as important predictors within the physicochemical-descriptor model. Plotting hydrophobicity against isoelectric point revealed partial separation of the high- and low-FRET subpopulations. To place these differences in context, we also plotted the mean values of representative sets of disordered, coiled-coil, globular and transmembrane proteins (Figure 3g).

These analyses show that FRET-defined phenotypes are predictable from sequence within the sampled random library. Global physicochemical descriptors provided substantial predictive information, but the strongest classification performance was extracted from more complex models that learn directly from the sequence.

## Discussion

A central question in evolutionary biology and protein biophysics is to what extent the structural diversity observed in nature reflects the intrinsic physical properties of polypeptides and how these properties have been shaped by evolutionary constraints ^27^ (Chan and Bornberg-Bauer, 2002). Theoretical work has proposed that elements of protein structure can arise from general physical properties of polypeptide chains^28–30^, while experimental sampling has identified random sequences with secondary structure and proteolytic resistance ^10,18,31^. At the same time, random sequences with higher structural content have shown lower solubility than sequences with high predicted disorder. In natural globular proteins, aggregation-prone regions contribute to thermodynamic stability. Furthermore, folded protein complexes can acquire the capacity for supramolecular assembly through only a few surface mutations, without losing their native folds ^32,33^. These observations motivate a distinction between the accessibility of structural tendencies and the emergence of sequences that combine stable, well-defined conformations with solubility.

In this study, we systematically explored the conformational and assembly properties of a large random sequence library. Previous efforts to probe such spaces have measured solubility and proteolytic resistance or quantified amyloid nucleation ^9,10^. Using high-throughput FRET-FACS selection calibrated with fluorescence and anisotropy imaging measurements of control proteins, we enriched populations with contrasting FRET signals and distinguished different behaviors among their members. Low-FRET candidates included both diffusely distributed proteins (resembling the soluble, disordered control), and localized assemblies. High-FRET candidates displayed IB-associated fluorescence, but several also showed substantial FRET in cytosolic regions outside these assemblies, with different FRET efficiencies in the two locations. Comparison with controls characterized by fluorescence anisotropy suggests that the contrasting assembly-associated signals reflect differences in local reporter packing, with closer intermolecular proximity contributing to high FRET. Assembly formation was also not accompanied by a uniform chaperone-associated response: several high-FRET candidates, particularly those recovered in H3, showed relatively little *ibpA* activation. Thus, even variants that formed visible assemblies differed in the signals associated with their packing and the cellular responses they elicited. These differences would be obscured by grouping all variants as simply soluble or aggregated.

The sorted populations provide starting material for investigating whether structural tendencies can coexist with solubility and stability in individual sequences. The cytosolic FRET detected in several candidates raises the possibility that compact conformations are accessible even without evolutionary optimization and for sequences that also form larger assemblies. However, across successive high FRET sorting rounds, the soluble fraction decreased when the proteins were expressed without the fluorescent reporters. Continued enrichment for high FRET may therefore increasingly favor intermolecular contributions to the signal. H1 and H2 thus offer particularly useful starting populations for secondary screening, supported both by cytosolic FRET in individual candidates and by their greater (reporter-free) soluble population relative to H3. The next question is whether the cytosolic signal arises from compact monomers, soluble oligomers or more heterogeneous conformations. Secondary screening for solubility and stability, followed by characterization of individual proteins could distinguish these possibilities.

If some degree of structural organization can arise before extensive sequence optimization, an important next question is how such properties might contribute to the earliest stages of protein evolution. A newly translated sequence does not need to resemble a modern protein immediately to become evolutionarily relevant; it first needs to be sufficiently compatible with the cellular environment to persist, while offering some physical property on which selection can act. This problem is particularly relevant to *de novo* gene birth, where new proteins emerge from previously non-coding DNA and therefore begin with little or no history of selection ^34,35^. Functional proteins have already been recovered from random sequence libraries, most often through binding or interactions with existing cellular components, indicating that useful molecular interactions can arise from largely unevolved sequences ^7,36–38^. The emergence of natural *de novo* proteins likewise implies that at least some newly translated sequences can avoid immediate elimination and acquire selectable effects ^39–41^. Could some of these sequences also adopt soluble, compact conformations that make them particularly suitable starting points for further evolution? One possibility is that partially organized conformations provide “proto-folds”, i.e. structures that need not possess the stability or specificity of established proteins but could support interactions or rudimentary activities on which selection can act ^42^. Evolution could then refine these tendencies by stabilizing productive conformations, limiting detrimental association and improving functional specificity. In this view, the key evolutionary step may not be the *de novo* appearance of a fully optimized fold, but the progressive enrichment of sequence features that make a nascent protein more soluble, stable and functionally exploitable.

An important implication of our results is that these experimentally defined phenotypes are not only selectable, but also learnable from sequence. Classifiers trained on the FRET-defined populations distinguished low- and high-FRET variants within the sampled library, while simpler physicochemical descriptors captured part, but not all, of this signal. This suggests that iterative experimental selection could be coupled to sequence-based learning to progressively enrich combinations of properties that are difficult to isolate through a single readout. In the context of the secondary screening proposed above, solubility, stability and reporter-free structural behavior could provide additional labels that refine the models beyond FRET alone. Such an approach would turn the screen from a one-step classification of random sequence space into an iterative search for sequence features that move unevolved proteins towards soluble, stable and potentially functional states.

Several limitations define the present interpretation. FRET reports fluorophore proximity and can contain both intra- and intermolecular contributions, while the fluorescent fusions themselves may influence the behavior of the inserted sequences. Our individual characterization also focused on fusion constructs, while reporter-free solubility was measured at the population level. We therefore cannot yet determine how frequently the selected populations contain compact monomeric proteins, nor whether cytosolic FRET reflects monomers, soluble oligomers or conformationally heterogeneous species. At the same time, structural predictions do not currently resolve this uncertainty. The low confidence of models generated for these sequences limits their use for assigning specific folds, particularly in regions of sequence space poorly represented in evolutionary training data ^43^. Finally, the 20F109 library samples one protein length and one amino acid distribution, and therefore represents only a small, compositionally defined region of the vastly larger random sequence space. These limitations motivate deeper sequence-resolved characterization rather than changing the central observation that distinct conformational and assembly-associated phenotypes can be experimentally enriched and learned from sequence.

The protein universe accessible to biology is only a small evolutionary sample of what polypeptide chemistry permits. By experimentally probing sequence space outside this sample, our study begins to resolve the “negative space” surrounding natural proteins - sequences that have not been optimized by evolution, yet already display distinguishable and learnable physical behaviors. Some of these behaviors lead to disordered states or aggregation, while others may provide starting points from which solubility, stability and eventually function can be refined. Determining how far such starting points lie from useful protein solutions is central both to understanding how new proteins can emerge from previously non-coding sequences and to learning which properties evolution must actually build, rather than merely improve. In turn, experimentally mapping this unevolved sequence space could provide protein design with information fundamentally different from that contained in natural sequence and structure databases. Instead of learning only from solutions already discovered by biology, future design approaches could also learn from the routes leading toward (and away from) those and other solutions.

## Materials and methods

### Library design and synthesis

The coding sequence of the library was generated by the CoLiDe tool ^18^. The parameters were set to generate 107 positions using degenerate codons with a maximum ratio of an amino acid in a single codon of 0.9. After obtaining the CoLiDe solution, two additional codons were introduced after codon position 53 to serve as a ligation point in subsequent library assembly. The flanking positions in this sextet were degenerated to maintain variability of the codons, while the central four nucleotides served as 4 base pair overhang in subsequent restriction cloning. Next, the string was split in half and cloning elements were added. Both termini of the sequence were furnished with short handles for primer annealing, BsaI enzyme recognition site, and 4 base pair overhang sequence. The positive strand of the coding sequences were ordered as PAGE Ultramer oligos at a nanomolar scale from IDT(Integrated DNA Technologies, Coralville, IA).

The library was assembled from two 199-nucleotide-long oligos. To synthesize the second strand, we used a primer extension with the Large (Klenow) fragment of DNA polymerase I (New England Biolabs, M0210L). Reaction was mixed as follows: First, 1.55 nanomole of the library oligo was mixed with 1.55 nanomole of the primer, NEBuffer™ 2 buffer and water, heated to 95 °C, cooled to 25 °C at 0.1 °C/s pace (with 5 min hold at 71 °C - primer T_m_). After reaching 25 °C, the polymerase and dNTPs were added to the reaction and incubated for 90 minutes. The primer melting temperature was calculated according to the approximate composition of 1X NEBuffer™ 2. The mixture was purified with the Monarch PCR clean-up kit (New England Biolabs, T1030) and the concentration was measured with Nanodrop spectrophotometer (Thermo Fisher Scientific Inc., Waltham, MA).

### Library assembly and cloning

Double-stranded products of Klenow extension reaction were assembled with the recipient vector pETMF using the NEBridge® Golden Gate Assembly Kit (New England Biolabs, E1601S). Details of pETMF construction are described in the Aubel et al. study ^12^. In short, the vector allows T7-controlled expression of the insert protein fused to a genetically encoded FRET pair (mTurquoise2 and mVenus). 60 fmol of pETMF was mixed with 120 fmol of both dsDNA library fragments in 60 µl reaction and incubated in 5 min cycles of 16 °C and 37 °C for 16 hours. The reaction was then purified with the Monarch PCR clean-up kit and eluted to 10 µl. Next, 2 µl of purified ligation mixture was electroporated into 25 µl E. cloni 10G Elite (LGC Inc., 60052-1) in two rounds, each batch was recovered in 975 µl of recovery media and pooled. After 1 hour of recovery growth at 37 °C shaking, 1950 µl of culture was plated on a single LB-kanamycin 245 mm Bioassay plate and incubated at 37 °C for 16 hours. The remaining 50 µl of recovered cells were used for serial dilution and plated on LB-kanamycin dishes to estimate the efficiency of transformation. Following the incubation, the plate was washed with cold LB-kanamycin and the colonies were carefully scraped. An appropriate amount of cells (as measured by OD_600_) were used for plasmid isolation with the Zymo Midiprep kit. Colonies from dilution plates were used to check the insert integrity by colony PCR. The pooled library prep was used to transform E. coli expression strain. Namely, 25 µl of electrocompetent E. cloni EXPRESS (LGC Inc., 60401-2) cells were electroporated with 50 ng of pETMF-20F109 plasmid pool and again plated on a 245 mm Bioassay dish, followed by incubation at 30 °C for 16 hours to prevent overgrowth. Analogically, the plates were washed with cold LB-kanamycin, colonies scraped, and the optical density of the culture measured. Cells were used for both plasmid DNA preparation and the generation of glycerol stocks. The cultures were diluted to a final OD_600_ = 1, glycerol concentration 20%, and stored in -80 °C.

### Protein expression and purification

We used *E. coli* BL21(DE3) expression strain (E. cloni EXPRESS, LGC Inc., 60401-2) for FACS, flow cytometry, fluorescence spectroscopy, and microscopy experiments. Overnight cultures were inoculated from glycerol stocks and incubated for 16 hours at 37 °C in LB media supplemented with 50 µg/ml of kanamycin and 1 % glucose. Next, the cells were transferred to fresh LB-kanamycin and the expression was induced during mid-log phase by 0.5 mM IPTG and incubated shaking at 220 rpm for 18 hours at 25 °C in LB-kanamycin media. To prevent genetic drift, the cells used for the FACS were cultured in 50 ml of LB-media. For protein purification the expression was scaled up to 500 ml. Following expression at 25 °C for 18 hours, the cells were harvested (12 000 x G, 4 °C, 10 min), the pellets were weighted and resuspended in lysis buffer (1x BugBuster, 50 mM sodium phosphate, 300 mM sodium chloride, 0.1 % Triton X-100, 1x cOmplete protease inhibitor, 12 U/ml benzonase, 1 mg/ml lysozyme, pH = 6.8). After 30 min incubation with gentle agitation, the lysate was passed through glass homogenizer and the insoluble fraction was separated by centrifugation at 30 000 x G, 4 °C for 30 minutes. The supernatant was loaded on a gravity-flow IMAC column (TALON® Metal Affinity Resin, Shiga, JP) pre-equilibrated with buffer A (50 mM sodium phosphate, 300 mM sodium chloride, 0.1 % Triton X-100, 1x cOmplete protease inhibitor, pH = 6.8) and incubated with the resin on a tumbler at 4 °C for 45 minutes. The column was washed twice with 10 bed volumes of buffer B (50 mM sodium phosphate, 300 mM sodium chloride, 1x cOmplete protease inhibitor, pH = 6.8) and the Hig-tagged proteins were finally eluted with 5 bed volumes of buffer C (50 mM sodium phosphate, 300 mM sodium chloride, 300 mM imidazole, 0.05 % Triton X-100, 1x cOmplete protease inhibitor, pH = 6.8). Next, the eluted fraction were concentrated in 15 ml Amicon® Ultra Centrifugal Filter with 10 kDa cutoff, loaded to HiPrep 16/60 Sephacryl S-300 HR column equilibrated with buffer B and run at 0.5 ml/min flow rate. The peaks were collected manually based on 220 nm and 280 nm absorbance, the collected samples were further concentrated in 10 kDa Amicon® Ultra Centrifugal Filter and the concentration was assessed by absorbance at 280 nm or using Bradford assay, before flash freezing in liquid nitrogen.

### Spectroscopy data acquisition and analysis

To assess the FRET efficiencies we measured donor lifetime by time-correlated single-photon counting on cell lysates for general screening and on purified samples for the selected controls. In cell-lysate experiments, *E. coli* cultures were diluted to OD_600_ = 1 and harvested (4000 x G, 4 °C, 5 min). The pellets were resuspended in 1x BugBuster® Master Mix (Milipore, 71456-3), incubated at room temperature with gentle shaking for 30 minutes and the lysates were clarified by centrifugation (20 000 x G, 4 °C, 20 min). Purified samples were thawed and centrifuged. Supernatants were diluted to A_405_ = 0.1 to prevent inner filter effect. The measurements were carried out at room temperature in quartz glass cuvettes (Hellma®, Sigma-Aldrich, Z600393-1EA) on Photoluminescence Spectrometer FLS 1000 (Edinburgh Instruments Ltd., Livingston, UK) equipped with Xenon lamp for steady-state measurements and 405 nm picosecond pulsed diode laser (EPL-405, Edinburgh Instruments Ltd., Livingston, UK) for TCSPC. We collected emission spectra of the donor (mTurquiose2) in cell-lysate with 405 nm excitation and determined the emission maximum at 474 nm. TCSPC measurements were recorded for 474 nm emission, with slits at 0.5 nm and 10 MHz laser repetition rate. We recorded 10 000 events in 50 ns window with 2048 channels. To measure the instrument response function (IRF), we used 0.1 % solution of colloidal silica (LUDOX^®^ TM-50, Sigma-Aldrich, 420778-1L).

### Analysis of fluorescence decay curves

Time-resolved fluorescence traces (emitted photons as a function of time) were subjected to lifetime distribution analysis by the maximum entropy method as previously described ^44–46^ (Lórenz-Fonfría and Kandori 2006; Lórenz-Fonfría and Kandori 2007; Lórenz-Fonfría and Kandori 2007). To this end, the datasets were numerically transformed from the time domain to the lifetime domain using the inverse Laplace transform with the aim of finding the lifetime distribution that maximizes an entropy function while minimizing the squared sum of the residuals between the data and the fit. The time constants (τ) were fixed and logarithmically distributed from 0.01 ns to 100 ns with 41 points per decade. Thus, the amplitudes (A) were the only fitting parameters.

For each peak *i* in the lifetime distribution (typically containing 2 peaks, see Fig. S1), the mean lifetimes τ_i_ (amplitude-weighted) were retrieved:

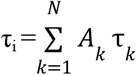

Where *A_k_* are the normalized amplitudes across the peak 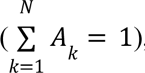, and τ*_k_* represents the individual lifetimes constituting the peak.

Similarly, the mean lifetime (amplitude-weighted) of the whole distribution was calculated as:

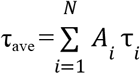

*A_i_* are the normalized amplitudes of the i-th component 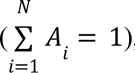, and τ*_i_* represents the lifetime of the i-th component.

FRET efficiencies were calculated from the mean lifetimes of the full distribution:

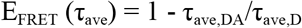

Where DA is the donor-acceptor sample (1259, 1856, 665, etc.) and D is the donor-only sample.

### Image processing and FLIM data analysis

Image processing and FLIM analysis were done in Python with custom Jupyter notebooks, available at [repository URL]. Time-tagged photon data (PicoQuant .ptu) were reconstructed with tttrkit (https://github.com/IMCF-Biocev/tttrkit). Segmentation used scikit-image and Cellpose-SAM, and masks were curated in napari.

### Intensity images and segmentation

After correcting time-tag overflows, photons were assigned to pixels using the line and frame markers. The scan geometry was read separately for each file from its .ptu header and the matching.msr file. Intensity images were computed as photon counts summed over both detection channels (mTurquoise2 and mVenus) and all z-planes. Inclusion bodies (IBs) were segmented from these images with a difference-of-Gaussians filter and Otsu thresholding. Cells were segmented with Cellpose-SAM. Where a transmitted-light z-stack had been recorded alongside the FLIM data, Cellpose was given its per-pixel standard deviation across z-planes. Otherwise, Cellpose was given the fluorescence intensity image with the IBs clipped, so that they did not dominate segmentation. Cytosolic masks were defined as cell masks without the IBs and their immediate surroundings. All masks were then inspected and curated manually in napari, and images unsuitable for analysis were discarded.

### Extraction of decays and phasors

Lifetime analysis used only photons from the donor (mTurquoise2) detection channel, with all z-planes summed. For every pixel within the masks, the photon count and the first-harmonic phasor at the laser repetition frequency (f = 40 MHz, ω = 2πf), g = ⟨cos ωt⟩ and s = ⟨sin ωt⟩, were accumulated directly from the photon arrival times t at the native TCSPC resolution. Phasors of single objects (one IB or cytosolic region), images and constructs were calculated as photon-weighted averages of the per-pixel phasors, which is equal to the phasor of the summed decay. Objects with fewer than 300 photons were excluded.

### Day-to-day correction and phasor calibration

Drift of the instrument timing between imaging sessions was corrected from the rising edge of the accumulated IB decays: the phasors of each session were rotated to match the shift of the rising edge relative to the reference session. The phasor plane was then calibrated against a donor-only (mTurquoise2) sample measured in the reference session. The complex calibration factor c = Φ_theor_/Φ_meas_ was calculated from the measured phasor of its cytosolic signal, Φ_meas_, and the theoretical single-exponential phasor, Φ_theor_ = 1/(1 − iωτ_D_), with the literature lifetime τ_D_ = 3.8 ns ^47^. This factor was applied to all phasors, combined with the session rotation. The phase lifetime was calculated as τ_φ_ = s/(ωg) and the apparent FRET efficiency as *E*_φ_ = 1 − τ_φ_/τ_D_. In the phasor plots, each construct is shown as one pooled IB phasor and one pooled cytosolic phasor, connected by an arrow from cytosol to IB.

### Intensity-modulated lifetime maps

Lifetime maps were generated from the calibrated per-pixel donor phasors. To reduce shot noise, phasors and photon counts were summed over a 3 × 3 px sliding window before τφ was calculated. Hue encodes τφ on an isoluminant colour map (constant CIE L*), so that it stays independent of brightness, which encodes the pooled photon count. All maps share one lifetime scale, set by the 2nd–98th photon-weighted percentile of pixels with at least 30 photons. For brightness, photon counts were percentile-stretched for each image and a gamma of 0.7 was applied.

### Simultaneous Homo- and Hetero-FRET analysis

Simultaneous Time-resolved anisotropy of acceptor and FLIM-FRET of donor analysis was performed on Leica SP8 TCS SMD, using 63x/1.2 NA water objective, 1 AU pinhole, HyD detection range 450-495 nm for donor and 515-600 nm for acceptor, 512x512 pixels, 37 nm pixel size, line accumulation 16, 600 Hz bidirectional scan, 10 consecutive time points to mitigate the effect of sample drift and bleaching. Each timepoint consisted of three frames, first and second frame with linearly polarized 505 nm WLL excitation and perpendicular and parallel, respectively, detection, and the third frame with 405 nm laser diode excitation. Both lasers were synchronized at 20 MHz repetition frequency. Photon detection events from both HyD detectors were recorded with MultiHarp 150 (SPT64 software, PicoQuant, Germany) with 40 ps sampling using TTTR data format. FLIM image reconstruction including ROI application and time-resolved anisotropy fitting was done in custom TTTR Data Analysis software (developed by Aleš Benda, written in LabVIEW).

For time resolved anisotropy the G-factor (1,159) for the used setup was estimated from measurement of fast rotating Atto488 in water (rotational correlation time ∼290 ps in water at 37°C). The fluorescence anisotropy decay was calculated from the histograms for parallel and perpendicular signal using the standard formula, including correction for constant background:

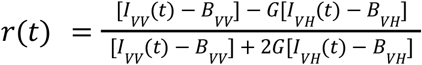

The standard deviation for each time channel of the anisotropy decay curve was estimated as the standard error for sliding 11 time channels window (5 previous time channels, the given time channel and next 5 time channels). The weighted tail-fit using non-linear least square approach started 160 ps after IRF maximum to avoid contribution of reflected and scattered light. Two-exponential fit was applied to all decays with fluorescence proteins, with fast ns component reflecting homo-FRET and slow tens of ns component reflecting the rotation of the fluorescent proteins.

For steady-state anisotropy calculation only photons in time channels starting 160 ps after IRF maximum were summed up and corrected for constant background.

The hetero-FRET between the donor and the acceptor was quantified by weighted tail-fit of donor only fluorescence histograms, using non-linear least square approach starting 160 ps after IRF maximum to avoid contribution of reflected and scattered light. Two exponentials were used to fit the histograms to obtain reasonably good and robust fits and amplitude-weighted average lifetime was calculated as a quantification of the hetero-FRET.

### Fluorescence lifetime imaging microscopy

To assess the localization and fluorescence lifetime *in vivo*, FLIM data were collected. The cells were prepared as described in the *Protein expression and purification* section, washed with cold PBS and immobilized on poly-L-lysine coated 18-well µ-Slide (Ibidi). Imaging was performed on an Abberior Infinity STED microscope using an Olympus UPLXAPO 60× oil immersion objective (NA 1.42). Donor excitation was achieved with a 440 nm pulsed laser (40 MHz repetition rate). Emission was collected at 460–510 nm (donor) and 520–570 nm (acceptor). To minimize focusing-related variability, z-stacks of five planes were acquired per field of view with 0.6 µm spacing and 60 nm pixel size. Raw data were processed using a custom Python pipeline.

### Flow cytometry and library sorting

The expressing *E. coli* cultures used for fluorescence activated cell sorting (FACS) were initiated from frozen glycerol stocks (prepared as in *Library assembly and cloning*). The day before sorting, the stocks were thawed and diluted 5 times with LB + kanamycine to OD_600_ = 0.2 incubate in a shaker at 37 °C until OD_600_ = 0.6 (typically 60 minutes). The cultures cooled down, induced 0.5 mM IPTG and incubated in shaker at 25 °C for 16 hours. The cells were washed three times with cold PBS, diluted to OD_600_ = 0.03 and analysed on either BD LSRFortessa or BD FACSAria Fusion. Three fluorescence channels were used: donor channel with 405 nm excitation and 450/50 bandpass filter for emission; FRET channel with 405 nm excitation and 540/40 filter for emission; acceptor channel with 488 nm excitation and 530/30 filter for emission. Following the shape and size gating, the events positive for both donor and acceptor fluorescence were gated (P1 gate). Next, a parameter FRET ratio was derived from the intensity of the FRET channel divided by donor channel intensity and the P1 population was projected as a histogram of the FRET ratio. To separate the low and the high FRET populations, the bottom and the top 10 % of the FRET ratio distribution were gated in the first round of the sorting, for any further round only the population in corresponding direction was sorted. For the sort report see *Extended data.* The cells were sorted to SOC media, recovered on LB-agar plates and incubated at 30 °C for 18 hours. The colonies were processed as in *Library assembly and cloning*.

### Next-generation sequencing

The library sequences were PCR amplified from the pETMF plasmid pools and internal barcodes were introduced. Quality control and NGS sequencing libraries preparation is described in detail in Aubel et al. ^12^. Pooled sequencing library was analysed on NextSeq platform (Illumina).

The raw reads from NGS analysis were filtered, quality checked and merged with fastp suit ^48^. Cutadapt was used to demultiplex the pooled samples ^49^. To generate a reference sequence database, the naive library pool was length filtered, flanking sequences were trimmed with cutadapt and redundancy was removed by clustering at 99% identity with cd-hit ^50^. Correct length open reading frames were extracted with custom Python script and translated to protein sequences. To obtain the counts of the variants in FACS sorted populations, the demultiplexed reads were mapped to the nucleotide reference library using Salmon ^51^.

### Cell free protein expression

The samples for solubility assessment and Lon protease resistance assay were expressed in the PUREfrex® 2.0 system (GeneFrontier, Chiba, JP) supplied with mRNA template. To prepare the mRNA, coding sequences were PCR amplified from the library pools and subcloned to pET-24a(+) with the cloning site modified to contain BsaI sites and FLAG sequence on the C-terminus of the open reading frame. The BsaI sites were introduced to the insert sequences by PCR, along with scars resulting from primer handles (N-terminal MLGGS, C-terminal SGGSELTS). The insert and the vector DNA were assembled in Golden Gate reaction and transformed to electrocompetent E. coli 10G cells and the plasmid pools were isolated from scrapped colonies. To generate the mRNA, the expression cassette was first PCR amplified from the plasmid pools with standard T7 sequencing primers (T7prom and T7term), as we found linear PCR fragments to be the optimal template for *in vitro* transcription. The *in vitro* transcription was carried out using HiScribe® T7 Quick High Yield RNA Synthesis Kit (New England Biolabs, E2050L) according to manufacturer recommendations and the RNA was purified by lithium chloride precipitation. Cell free protein synthesis was initiated by addition of template RNA to final concentration 0.3 µg/µl and supplemented with 0.05 % Triton X-100. After 3 hours incubation at 37 °C, the reaction was quenched by 5x dilution with puromycin buffer (10 mM HEPES, 300 mM NaCl, 0.05 % Triton X-100, 10 µg/ml puromycin, pH = 7.4). To separate soluble fractions, the samples were centrifuged at 21 000 x G for 30 minutes at 20 °C and analyzed by dot-blot. 2 µl of total and soluble fraction were loaded on nitrocellulose membrane and let air dry for 45 minutes. The membrane was blocked by 3 % BSA in TBST in 50 ml falcon tube on a roller for 1 hour, then incubated for 1 hour with 1:10000 dilution of ANTI-FLAG® M2-Peroxidase (HRP) antibody in 1 % BSA in TBST. The membranes were washed 3 times with TBS and incubated with Immobilon® Forte HRP substrate prior visualization.

### Chaperone response assay

Fluorescent protein under control of *ibpA* promoter was used to monitor the cellular heat-shock machinery response to expression of selected library variants. pSEVA631 was modified to carry ampicillin resistance and rigged with *ibpA-GFP* cassette. The reporter plasmid was co-transformed with pETMF plasmid carrying selected library variants and control proteins to *E. coli* BL21(DE3). The cells were cultured in deep well 96-well plates according to standard procedure: overnight growth in plate incubator at 37 °C and 1000 rpm in LB-kanamycin supplemented with 100 µg/ml ampicillin and 1 % glucose; re-inoculation to fresh LB-Kan-Amp media, growth at 37 °C and 1000 rpm until mid-log phase and induction of expression by 0.5 mM IPTG. Following 1 hour of expression at 37 °C the cells were cooled down and diluted 10-fold with PBS in black 96-well plates (Thermo Scientific™, Catalog no. 137101). The fluorescence was measured on Infinite M Plex microplate reader (Tecan Group Ltd., CHE) with following settings: GFP was excited with 465/9 nm and emission was detected in 510/20 nm bandwidth; YFP was excited with 516/9 nm and emission was detected in 545/20 bandwidth. The cell cultures were prepared in 3 independent replicates.

### Predictions

#### 3D structure predictions and derived metrics

3D structures of the individual random proteins were predicted by ESMFold (esm.pretrained.esmfold_v1)^2^ and with ColabFold (v1.6.1; colabfold_batch) using single-sequence MSA mode, one recycling iteration, one model, and random seed 0 ^52^. Secondary structure (SS) elements were extracted with the DSSP algorithm (v4.4.0) ^53^. Following types of SS were considered as structured fraction of the proteins: H = α-helix, E = extended strand, participates in β ladder, I = π-helix, G = 3_10_-helix, P = κ-helix (poly-proline II helix). For the more detailed plots: the counted elements for helix were H, G and P, for strand only E and for turn and bend T = hydrogen-bonded turn and S = bend, respectively. For loops, the counted elements were B = residue in isolated β-bridge, T and S. Counted types of the extracted SS type were divided by protein length to get a fraction of the selected type for the whole protein.

NtoC termini distance was calculated with the formula:

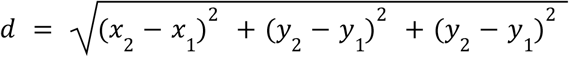

where *d* is the euclidean distance, *x_1_*, *y_1_*and *z_1_* are N termini C-alpha atom coordinates and *x_2_*, *y_2_* and *z_2_* are C-alpha atom coordinates.

The Radius of gyration was calculated with the formula:

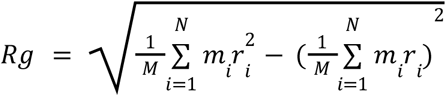

where M is the total mass of the protein, m_i_ is the mass of the i-th atom, r_i_ is the position vector of the i-th atom, and the masses used for individual atoms: C = 12.0107, O = 15.9994, N = 14.0067.

Solvent accessible surface area (SASA) for the whole protein was calculated by summing all solvent accessibility predicted for every AA. The theoretical isoelectric point was calculated using the Biopython library. The GRAVY score was calculated based on the Kyte-Doolittle amino acid hydrophobicity scale.

### Data Filtering & Dataset selection

#### Filtering steps

Populations in Figure 2a and Supplementary Figure 2b were filtered based on NGS read counts with following logic: percentile based filters were set to cover ∼80 % of total read count for corresponding population, therefore L2 - 65th, L1 - 60th, H1 - 70th, H2 - 80th and H3 - 90th percentile. For training dataset selection, first all sequences that were present in at least one of the performed sorting rounds were picked (694,032 sequences in total). To divide sequences into two subsets and remove duplicates, the relative counts of sequences from two rounds of low-FRET signal and three rounds of high-FRET signal were summed. Then these sums were compared to classify sequences: those with higher low-FRET summed counts were labeled as low-FRET, and vice versa. To further minimize false positives and negatives, the relative difference between the two sums was calculated and any sequence where this difference fell below a threshold of 0.1 was discarded, indicating that the counts in both subsets were too similar for reliable classification. After this filtering step, the dataset consisted of 238,635 sequences in the high-FRET subset and 343,457 sequences in the low-FRET subset. Following this filtering step, all sequences from all three rounds that had summed counts higher than 10 were selected, which resulted in 51,770 sequences labeled as low-FRET and 42,230 sequences labeled as high-FRET. This was the initial input into the further steps of the computational pipeline.

#### MMseqs2-guided dataset split

The train/test split was generated using a custom MMseqs2-based pipeline ^25^. All protein sequences were first clustered with MMseqs2 easy-cluster (minimum sequence identity 0.5, coverage 0.5, coverage mode 0, E-value 1×10⁻³). An all-against-all MMseqs2 search was then performed on the same sequences (createdb, search, convertalis; sensitivity 7.5, E-value 1×10⁻³, max-seqs 100). For each cluster, the mean sequence identity (seqid) of its members to sequences outside that cluster was computed from the search results (self-hits and within-cluster pairs excluded). Clusters were ranked by this external mean identity, and the 20% most distant clusters (lowest external mean seqid) were assigned entirely to the test set; all remaining clusters formed the training set. Because entire clusters were held out, no variant from a test cluster appeared in training, and train and test sets share no closely related variants within or across classes. In our dataset (94,000 variants), this yielded 38,497 clusters and a test set of 7,699 clusters containing 11,410 variants (12.1% of the dataset; 12.0% of high-FRET and 12.3% of low-FRET variants). Test-set selection was not performed independently within each class, and class balance in the test set was not explicitly enforced.

#### Data Splitting and Evaluation

For each run, except the MMseq2-guided split which was done beforehand and the dataset was held out as the test set, we used stratified random sampling to split the remaining data into training (80%) and modeling holdout (20%) sets, with a randomly generated seed saved for reproducibility. The training set was further stratified into optimization/training (80% of the training set) and validation (20% of the training set), yielding an effective 64/16/20 train/validation/modeling holdout partition. The validation subset was used for hyperparameter selection (when enabled), probability calibration via temperature scaling, and decision-threshold selection (maximizing F1). Final performance was then computed once on the modeling holdout using the fixed validation-derived calibration and threshold. This protocol was applied consistently across model architectures.

### Encoding for 2D-projection

#### PCA-Based 2D Projection

For low-dimensional visualization, two-dimensional PCA embeddings were generated for mean ESM, per-token ESM, and one-hot representations. Mean ESM and one-hot features were loaded in full, standardized (zero mean, unit variance), and projected with PCA (n_components=2).

#### UMAP-Based 2D Projection

Two-dimensional UMAP projections were generated for the same three encoding types. Mean ESM and one-hot matrices were projected directly with UMAP (n_components=2, fixed random state, low-memory mode). Cluster separation in the 2D projection was quantified using the silhouette coefficient and Davies–Bouldin index as implemented in scikit-learn (sklearn.metrics.silhouette_score and sklearn.metrics.davies_bouldin_score ^54^), computed on the 2D coordinates with experimental class labels as ground-truth groups. Higher silhouette and lower Davies–Bouldin indicate better cluster separation ^55,56^.

### Encoding for models

#### ESM-2 Embeddings

Protein sequences were encoded using pre-computed embeddings from ESM-2^2^, a transformer-based protein language model. Two types of embedding representations were extracted for each variant: (1) mean-pooled embeddings, representing the average of all per-token embeddings across the sequence (1280 dimensions), and (2) per-token embeddings, representing the full sequence-level representation obtained by flattening all token-level embeddings (111x1280, 142,080 when concatenated). Embeddings were extracted from layer 33 of the ESM-2 model (default for ESM-2-650M). For per-token embeddings, Principal Component Analysis (PCA) during data loading was applied to reduce dimensionality. All per-token embeddings were reduced from 142,080 dimensions to 2,560. This dimensionality reduction was performed before model training to reduce computational requirements while preserving the majority of variance in the embedding space.

#### One-Hot Encoding

Amino acid sequences were encoded using one-hot encoding, where each of the 20 canonical amino acids (A, C, D, E, F, G, H, I, K, L, M, N, P, Q, R, S, T, V, W, Y) was represented as a binary vector of length 20 with a single 1 at the position corresponding to that amino acid and zeros elsewhere. This resulted in a 3D feature matrix of shape (n_sequences, target_length: 111, 20), where each sequence was represented as a matrix of size (target_length: 111 × 20).

### Classification models

#### ESM-2 Embedding-Based Neural Network Models

For ESM-2 features, we trained separate models for mean and per-token embeddings using single-layer TensorFlow/Keras classifiers equivalent to L1-regularized logistic regression. Each model used a single dense layer with 2-unit softmax output and sparse categorical crossentropy loss, optimized with Adam. Input features were standardized with StandardScaler; for per-token embeddings, PCA reduction was applied for per-token embeddings. Hyperparameters (learning rate and L1 strength) were optimized with Optuna by maximizing validation accuracy (learning_rate: 1e-5-1e-1 log scale; l1_reg: 1e-6-1e-1 log scale; median pruner with startup/warmup settings), followed by final training with early stopping (patience=10).

#### One-Hot Encoding-Based Neural Network Model

One-hot encoded sequences were modeled with a two-hidden-layer MLP in TensorFlow/Keras: input (L, 20) (where L is the configured/potentially auto-adjusted target length), flattening, ReLU hidden layers, and a 2-unit softmax output. Default hidden-layer sizes were 64 and 32 units, and Optuna optimization searched n_units_l1 (16-256), n_units_l2 (8-128), batch_size (16/32/64/128), and epochs (20-100), with validation accuracy as the objective. The final model was then trained using the selected configuration on the training split with validation monitoring.

#### ESM-2 Embedding-Based Random Forest Models

For ESM-2 features, we trained separate RandomForestClassifier models for mean and per-token embeddings. Features were standardized prior to fitting; per-token representations were optionally PCA-reduced (same strategy as above). Hyperparameters were optimized with Optuna when enabled (n_estimators: 50-500, max_depth: 3-30, min_samples_split: 2-20, min_samples_leaf: 1-10, max_features: {sqrt, log2, None}); otherwise predefined defaults were used. Final models were fit with the selected settings (n_jobs=-1) and saved for reproducibility.

#### One-Hot Encoding-Based Random Forest Model

For one-hot features, encoded tensors (L, 20) were flattened to 1D vectors (L x 20) before training a RandomForestClassifier. Hyperparameters were tuned with Optuna using the same search space as above, or fixed defaults were used when optimization was disabled (n_estimators=200, max_depth=15, min_samples_split=5, min_samples_leaf=2, max_features=’sqrt’). Models were trained with class-stratified splits and run-level seeded randomness for reproducibility.

#### Sequence composition logistic regression baseline

As a composition-only reference model (no learned embeddings), we trained L2-regularized logistic regression in scikit-learn on hand-crafted sequence features from each variant FASTA sequence. Features comprised mean Kyte–Doolittle GRAVY, isoelectric point (pI) and net charge at pH 7 (Biopython), seven amino-acid group fractions defined as the proportion of residues in each physicochemical set—hydrophobic (L, V, I, G, A, P), basic (R, K, H), sulphur-containing (M, C), amide (Q, N), aromatic (F, W, Y), hydroxy (T, S), and acidic (D, E)—and Shannon entropy of amino-acid composition (11 features total). Input features were standardized with StandardScaler, and classification used LogisticRegression with L2 penalty, LBFGS solver, and max_iter=2000 (default C=1.0). Models were fit on the optimization split; on the validation split, probabilities were calibrated by temperature scaling (temperature minimizing expected calibration error over 100 log-spaced candidates from 0.01 to 10) and the decision threshold was set to maximize F1. Performance metrics, ROC curves, and permutation importance (ROC-AUC decrease, 30 shuffle repeats) were computed on the modeling holdout using the fixed validation-derived calibration and threshold. With ablation enabled, the same protocol was applied to single feature groups: hydrophobicity (GRAVY and hydrophobic-residue fraction jointly), pI, net charge, each remaining residue-class fraction alone, and sequence complexity.

#### Model’s evaluation metrics

Let *y_i_* ∈ {0, 1} be the true label, *p_i_* ∈ [0, 1] the predicted probability for the positive class. With confusion-matrix counts (*TP* (True Positives), *TN* (True Negatives), *FP* (False Positive), *FN* (False Negatives):

**Accuracy**

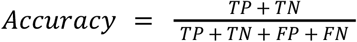

**Precision**

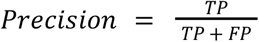

**Sensitivity**

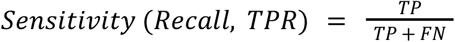

**F1 score**

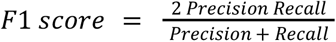

**Log Loss**

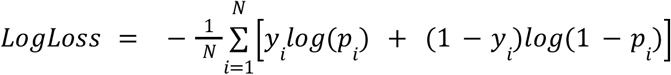

**AP**

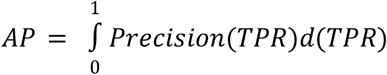

(implemented numerically from the precision-recall curve)

**ROC AUC**

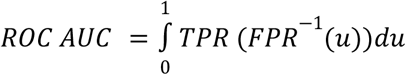

(implemented numerically from the ROC curve)

**Expected Calibration Error (ECE)**

It quantifies how far predicted probabilities are from observed outcome frequencies after binning predictions.

For *M* bins *B_m_* with *n_m_* = |*B_m_*|, confidence

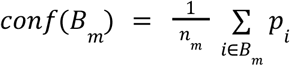

and accuracy

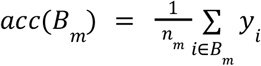

ECE is:

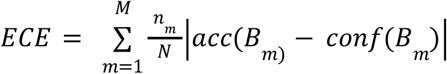

Lower ECE indicates better probability calibration.

ROC AUC, AP, log loss, and ECE are computed from probabilities and are threshold-independent (except ECE’s binning choice).

**Brier Score (BS)**

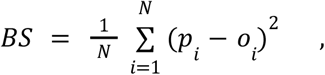

where *N* is total number of predictions tested, *p_i_* is the predicted probability high/low FRET class (a value between 0 and 1), *o_i_* is the actual sequence class

#### Threshold Optimization

Let *y_i_* ∈ {0, 1} be the true label, *p_i_* ∈ [0, 1] the predicted probability for the positive class, and 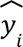 the predicted probability class after threshold optimization. A decision threshold *t* ∈ [0, 1] is applied to calibrated probabilities:

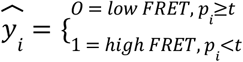

The threshold is selected on the validation set by scanning a grid of candidate values and choosing:

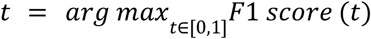

This fixed optimized *t* is then used for final test-set classification (MMseq2-guided split, hidden 100 most abundant sequences from each class, natural proteins classification).

#### Natural protein datasets

Natural sequences for embeddings plot in Figure S3c were obtained from SwissProt. 400 sequences from each kingdom with length 100-109 amino acids were filtered and mean embeddings were generated as described for the random protein variants. To compare the selected random proteins with natural proteins from diverse structural classes (Figure 4g), we generated following datasets: globular — α/◻ and α+◻ proteins with length range 85-300 amino acids from SCOPe 2.08 (code c and d); transmembrane — membrane and cell surface proteins in length range 85-300 amino acids from SCOPe 2.08 (code f); disordered — sequences from DisProt in length range 85-200; coiled coils — reviewed UniProt sequences with protein existence evidence, length 50-300 amino acids and annotated keyword coiled-coil (KW-0175).

**Figure 4.**
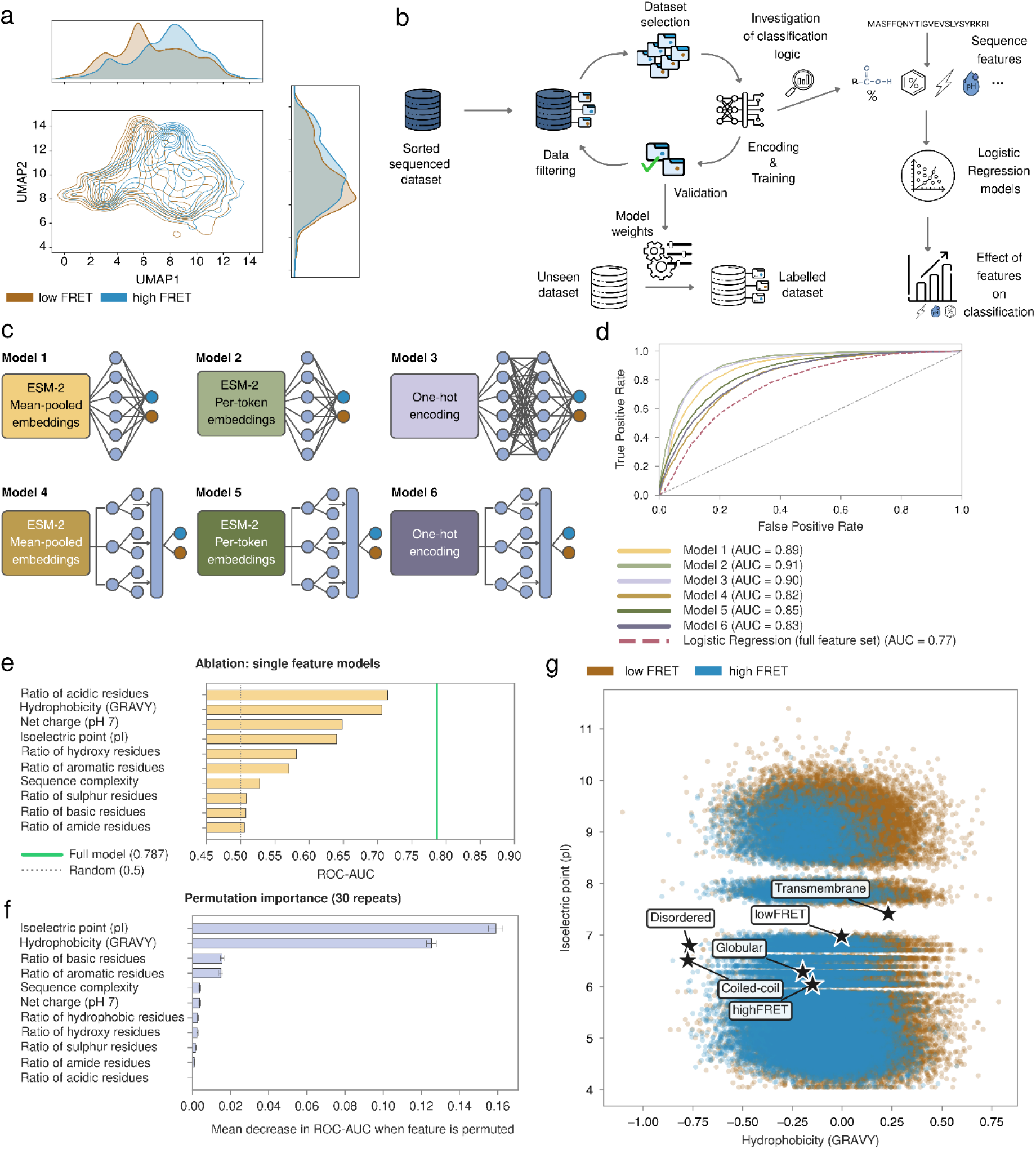
Overview of FRET Classification Models and Data Representation. **a**, Contour plot of 2D UMAP projected ESM-2 mean embeddings for the high/low-FRET dataset selected for supervised training of the classifiers. Colors indicate experimentally defined low-FRET (dark burgundy) and high-FRET (blue) populations from all rounds. **b,** Data handling pipeline for supervised classification of random sequences, illustrating the iterative selection of high- and low-FRET training data in the absence of clear class separation. Sequence features model serves to investigate classification logic. **c,** Architectures of the six trained classifiers. Neural Network type of models - *Model 1*: logistic regression using mean-pooled ESM-2 embeddings. *Model 2*: logistic regression using per-token ESM-2 embeddings. *Model 3*: multilayer perceptron (MLP) using one-hot encoded sequences as input and their Random Forest Classifier counterparts - *Model 4*, *Model 5* and *Model 6* with the same inputs in the respective order. All models output a probability (0-1); each sequence is classified as high-FRET if its calibrated probability is greater than or equal to the validation-optimized threshold chosen to maximize F1, and as low-FRET otherwise. **d,** Performance comparison of the six models on the held-out test dataset with usage of MMseqs2-guided diversity strategy, evaluated by ROC curves. **e,** Modeling-set ROC-AUC of logistic regression classifiers trained on one feature group at a time (ROC-AUC). Orange bars, individual feature-group models; green line, full 11-feature model (ROC-AUC = 0.787); grey dotted line, random performance (0.5). Hydrophobicity combines mean GRAVY and the fraction of hydrophobic residues (LVIGAP), since both measure the same compositional signal; all other bars use a single feature. **f,** Permutation importance (30 repeats). Mean decrease in ROC-AUC when each feature is randomly permuted in the full logistic regression model (error bars, standard deviation across repeats). Larger values indicate features that contribute more to classification in the multivariate model. **g,** Correlation plot of isoelectric point vs hydrophobicity (GRAVY). Each point is one library variant, coloured by FRET subpopulation (brown, low-FRET; blue, high-FRET). Stars mark mean GRAVY and pI for representative disordered (DisProt), coiled-coil (UniProt), globular, and transmembrane (SCOPe) protein sets, computed the same way as for the high/low-FRET subsets. The high-FRET centroid lies close to the globular reference, whereas the low-FRET centroid falls between the globular/disordered and transmembrane regions. Horizontal gaps in the point cloud (notably around pI ≈ 7 and 8) are library-design artefacts reflecting sequences excluded or under-represented by design (see Supplementary fig. 5e for reference library plot).

## Acknowledgments/funding

This work was funded by a research grant from the Volkswagen foundation (grant code 98183) and Primus grant (PRIMUS/20/SCI/012) from Charles University. The Institute of Biotechnology of the Czech Academy of Sciences acknowledges the institutional grant RVO86652036. We acknowledge CF Biophysics of CIISB, Instruct-CZ Centre BIOCEV, supported by MEYS CR (LM2023042) and ERDF-Project “UP CIISB” (No. CZ.02.1.01/0.0/0.0/18_046/0015974). Computational resources were provided by the e-INFRA CZ project (ID:90254), supported by the Ministry of Education, Youth and Sports of the Czech Republic. TN further acknowledges financial support from the Mobility Fund of Charles University and the IOCB Tech Foundation which supported her internship with MA. We would like to thank Theorema Inc., Tomáš Pluskal and Lars Eicholt for their consultation and their insightful comments on this project.

## Data and materials availability

Code used for imaging data analysis is available at Github repository https://github.com/buchelf-cuni/flim-fret-data-processing. Source data for cytometry, FACS, imaging and spectroscopy are deposited in https://zenodo.org/doi/10.5281/zenodo.23264574. All scripts used for model training, evaluation, ESM embedding extraction, and structure-based analyses, together with pre-trained classifiers and accompanying datasets, are publicly available on the Hugging Face Hub at https://huggingface.co/neuwirtt/FRET-FACS. The repository can be downloaded with hf download neuwirtt/FRET-FACS --local-dir FRET-FACS. Code is released under the MIT License; please cite this study when reusing the models or scripts.

## Notes

### Competing Interest Statement

The authors have declared no competing interest.

### Summary of Updates

Experimental results update: Phasor analysis and segmentation for FLIM data quantification. Fluorescence anisotropy results. Chaperone-response assay results updated with full control panel. Time-resolved spectroscopy on purified control proteins. Computational part updates: Binary classifiers revised. Single feature model and single feature permutation analysis added for interpretability.

## References

1. Jumper, J. et al. Highly accurate protein structure prediction with AlphaFold. Nature 596, 583–589 (2021).

2. Lin, Z. et al. Evolutionary-scale prediction of atomic-level protein structure with a language model. Science 379, 1123–1130 (2023).

3. Dauparas, J. et al. Robust deep learning–based protein sequence design using ProteinMPNN. Science 378, 49–56 (2022).

4. Ferruz, N., Schmidt, S. & Höcker, B. ProtGPT2 is a deep unsupervised language model for protein design. Nat. Commun. 13, 4348 (2022).

5. Ding, F. & Steinhardt, J. Protein language models are biased by unequal sequence sampling across the tree of life. 2024.03.07.584001 Preprint at 10.1101/2024.03.07.584001 (2024).

6. Keefe, A. D. & Szostak, J. W. Functional proteins from a random-sequence library. Nature 410, 715–718 (2001).

7. Frumkin, I. & Laub, M. T. Selection of a de novo gene that can promote survival of Escherichia coli by modulating protein homeostasis pathways. *Nat*. Ecol. Evol. 7, 2067–2079 (2023).

8. Schnettler, J. D. et al. Selection of a promiscuous minimalist cAMP phosphodiesterase from a library of de novo designed proteins. Nat. Chem. 16, 1200–1208 (2024).

9. Thompson, M. et al. Massive experimental quantification allows interpretable deep learning of protein aggregation. Sci. Adv. 11, eadt5111 (2025).

10. Tretyachenko, V. et al. Modern and prebiotic amino acids support distinct structural profiles in proteins. Open Biol. 12, 220040 (2022).

11. Philipps, B., Hennecke, J. & Glockshuber, R. FRET-based *in Vivo* Screening for Protein Folding and Increased Protein Stability. J. Mol. Biol. 327, 239–249 (2003).

12. Aubel, M. et al. High-throughput Selection of Human de novo-emerged sORFs with High Folding Potential. Genome Biol. Evol. 16, evae069 (2024).

13. Cheng, Z., Miskolzie, M. & Campbell, R. E. In Vivo Screening Identifies a Highly Folded β-Hairpin Peptide with a Structured Extension. ChemBioChem 8, 880–883 (2007).

14. Ebbinghaus, S., Dhar, A., McDonald, J. D. & Gruebele, M. Protein folding stability and dynamics imaged in a living cell. Nat. Methods 7, 319–323 (2010).

15. Shishido, H., Yoon, J. S. & Skach, W. R. A small molecule high throughput screening platform to profile conformational properties of nascent, ribosome-bound proteins. Sci. Rep. 12, 2509 (2022).

16. Wan, Q., Mouton, S. N., Veenhoff, L. M. & Boersma, A. J. A FRET-based method for monitoring structural transitions in protein self-organization. *Cell Rep*. Methods 2, (2022).

17. Tiwari, S., Fauvet, B., Assenza, S., De Los Rios, P. & Goloubinoff, P. A fluorescent multi-domain protein reveals the unfolding mechanism of Hsp70. Nat. Chem. Biol. 19, 198–205 (2023).

18. Tretyachenko, V. et al. Random protein sequences can form defined secondary structures and are well-tolerated in vivo. Sci. Rep. 7, 15449 (2017).

19. Evers, T. H., van Dongen, E. M. W. M., Faesen, A. C., Meijer, E. W. & Merkx, M. Quantitative Understanding of the Energy Transfer between Fluorescent Proteins Connected via Flexible Peptide Linkers. Biochemistry 45, 13183–13192 (2006).

20. Joshi, A. et al. Intermolecular energy migration via homoFRET captures the modulation in the material property of phase-separated biomolecular condensates. Nat. Commun. 15, 9215 (2024).

21. Tretyachenko, V., Voráček, V., Souček, R., Fujishima, K. & Hlouchová, K. CoLiDe: Combinatorial Library Design tool for probing protein sequence space. Bioinformatics 37, 482–489 (2021).

22. Schultz, T., Martinez, L. & de Marco, A. The evaluation of the factors that cause aggregation during recombinant expression in E. coli is simplified by the employment of an aggregation-sensitive reporter. Microb. Cell Factories 5, 28 (2006).

23. van Kempen, M. et al. Fast and accurate protein structure search with Foldseek. Nat. Biotechnol. 42, 243–246 (2024).

24. Lin, Z. et al. Evolutionary-scale prediction of atomic-level protein structure with a language model. Science 379, 1123–1130 (2023).

25. Steinegger, M. & Söding, J. MMseqs2 enables sensitive protein sequence searching for the analysis of massive data sets. Nat. Biotechnol. 35, 1026–1028 (2017).

26. Bushuiev, A., et al. Revealing data leakage in protein interaction benchmarks. Preprint at 10.48550/arXiv.2404.10457 (2024).

27. Chan, H. S. & Bornberg-Bauer, E. Perspectives on protein evolution from simple exact models. Appl. Bioinformatics 1, 121–144 (2002).

28. Ptitsyn, O. B. & Volkenstein, M. V. Protein Structures and Neutral Theory of Evolution. J. Biomol. Struct. Dyn. 4, 137–156 (1986).

29. Dill, K. A., Ozkan, S. B., Shell, M. S. & Weikl, T. R. The Protein Folding Problem. Annu. Rev. Biophys. 37, 289–316 (2008).

30. Kocher, C. D. & Dill, K. A. Origins of life: The Protein Folding Problem all over again? Proc. Natl. Acad. Sci. 121, e2315000121 (2024).

31. Chiarabelli, C. et al. Investigation of de novo Totally Random Biosequences, Part II. Chem. Biodivers. 3, 840–859 (2006).

32. Garcia-Seisdedos, H., Empereur-Mot, C., Elad, N. & Levy, E. D. Proteins evolve on the edge of supramolecular self-assembly. Nature 548, 244–247 (2017).

33. Romero-Romero, M. L. & Garcia-Seisdedos, H. Agglomeration: when folded proteins clump together. Biophys. Rev. 15, 1987–2003 (2023).

34. Bornberg-Bauer, E., Hlouchova, K. & Lange, A. Structure and function of naturally evolved *de novo* proteins. Curr. Opin. Struct. Biol. 68, 175–183 (2021).

35. Ruiz-Orera, J., Verdaguer-Grau, P., Villanueva-Cañas, J. L., Messeguer, X. & Albà, M. M. Translation of neutrally evolving peptides provides a basis for de novo gene evolution. *Nat*. Ecol. Evol. 2, 890–896 (2018).

36. Knopp, M. et al. A novel type of colistin resistance genes selected from random sequence space. PLOS Genet. 17, e1009227 (2021).

37. Neme, R., Amador, C., Yildirim, B., McConnell, E. & Tautz, D. Random sequences are an abundant source of bioactive RNAs or peptides. *Nat*. Ecol. Evol. 1, 0127 (2017).

38. Bhave, D. & Tautz, D. Effects of the Expression of Random Sequence Clones on Growth and Transcriptome Regulation in Escherichia coli. Genes 13, 53 (2022).

39. Vakirlis, N. et al. De novo emergence of adaptive membrane proteins from thymine-rich genomic sequences. Nat. Commun. 11, 781 (2020).

40. Vakirlis, N., Vance, Z., Duggan, K. M. & McLysaght, A. De novo birth of functional microproteins in the human lineage. Cell Rep. 41, (2022).

41. Chou, L. et al. Environmental stress and phenotypic tradeoff modulate the adaptive potential of novel coding sequences for de novo gene birth. 2026.08.24.746591 Preprint at 10.64898/2026.08.24.746591 (2026).

42. Papadopoulos, C. et al. Intergenic ORFs as elementary structural modules of de novo gene birth and protein evolution. Genome Res. 10.1101/gr.275638.121 (2021) doi:10.1101/gr.275638.121.

43. Middendorf, L. & Eicholt, L. A. Random, de novo, and conserved proteins: How structure and disorder predictors perform differently. Proteins Struct. Funct. Bioinforma. 92, 757–767 (2024).

44. Lórenz-Fonfría, V. A. & Kandori, H. Transformation of Time-Resolved Spectra to Lifetime-Resolved Spectra by Maximum Entropy Inversion of the Laplace Transform. Appl. Spectrosc. 60, 407–417 (2006).

45. Lórenz-Fonfría, V. A. & Kandori, H. Practical Aspects of the Maximum Entropy Inversion of the Laplace Transform for the Quantitative Analysis of Multi-Exponential Data. Appl. Spectrosc. 61, 74–84 (2007).

46. Lórenz-Fonfría, V. A. & Kandori, H. Bayesian Maximum Entropy (Two-Dimensional) Lifetime Distribution Reconstruction from Time-Resolved Spectroscopic Data. Appl. Spectrosc. 61, 428–443 (2007).

47. Goedhart, J. et al. Structure-guided evolution of cyan fluorescent proteins towards a quantum yield of 93%. Nat. Commun. 3, 751 (2012).

48. Chen, S., Zhou, Y., Chen, Y. & Gu, J. fastp: an ultra-fast all-in-one FASTQ preprocessor. Bioinformatics 34, i884–i890 (2018).

49. Martin, M. Cutadapt removes adapter sequences from high-throughput sequencing reads. EMBnet.journal 17, 10–12 (2011).

50. Li, W. & Godzik, A. Cd-hit: a fast program for clustering and comparing large sets of protein or nucleotide sequences. Bioinformatics 22, 1658–1659 (2006).

51. Patro, R., Duggal, G., Love, M. I., Irizarry, R. A. & Kingsford, C. Salmon: fast and bias-aware quantification of transcript expression using dual-phase inference. Nat. Methods 14, 417–419 (2017).

52. Mirdita, M. et al. ColabFold: making protein folding accessible to all. Nat. Methods 19, 679–682 (2022).

53. Hekkelman, M. L., Salmoral, D. Á., Perrakis, A. & Joosten, R. P. DSSP 4: FAIR annotation of protein secondary structure. Protein Sci. 34, e70208 (2025).

54. Pedregosa, F., et al. Scikit-learn: Machine Learning in Python. Preprint at 10.48550/arXiv.1201.0490 (2018).

55. Rousseeuw, P. J. Silhouettes: A graphical aid to the interpretation and validation of cluster analysis. J. Comput. Appl. Math. 20, 53–65 (1987).

56. Davies, D. L. & Bouldin, D. W. A Cluster Separation Measure. IEEE Trans. Pattern Anal. Mach. Intell. PAMI-1, 224–227 (1979).

