## Supplementary figures for "Learning the structural diversity in random protein sequence space"

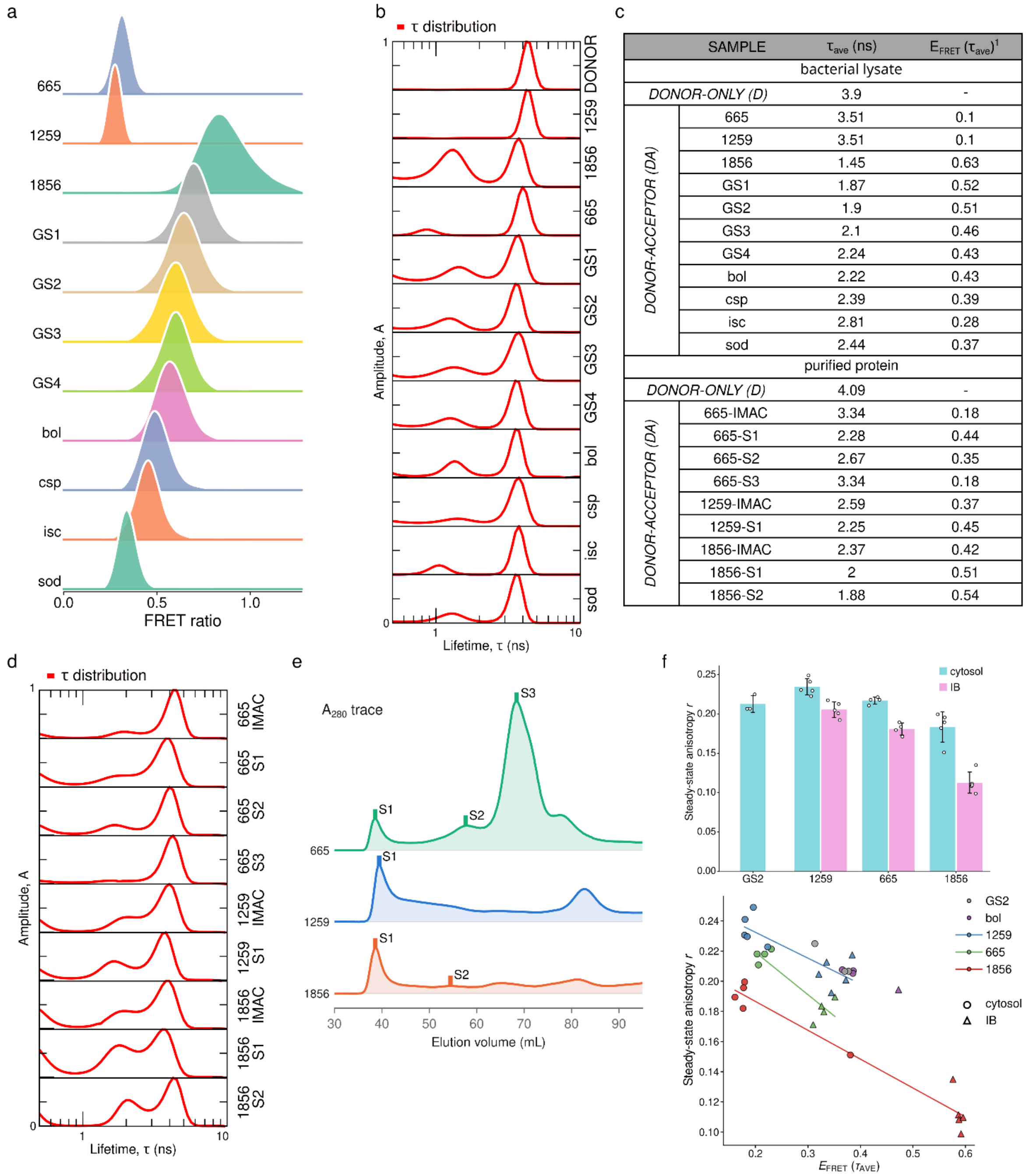


| SAMPLE | | $\tau_{ave}$ (ns) | $E_{FRET}$ ($\tau_{ave}$)[1] |
|---|---|---|---|
| bacterial lysate | | | |
| DONOR-ONLY (D) | | 3.9 | - |
| DONOR-ACCEPTOR (DA) | 665 | 3.51 | 0.1 |
| | 1259 | 3.51 | 0.1 |
| | 1856 | 1.45 | 0.63 |
| | GS1 | 1.87 | 0.52 |
| | GS2 | 1.9 | 0.51 |
| | GS3 | 2.1 | 0.46 |
| | GS4 | 2.24 | 0.43 |
| | bol | 2.22 | 0.43 |
| | csp | 2.39 | 0.39 |
| | isc | 2.81 | 0.28 |
| | sod | 2.44 | 0.37 |
| purified protein | | | |
| DONOR-ONLY (D) | | 4.09 | - |
| DONOR-ACCEPTOR (DA) | 665-IMAC | 3.34 | 0.18 |
| | 665-S1 | 2.28 | 0.44 |
| | 665-S2 | 2.67 | 0.35 |
| | 665-S3 | 3.34 | 0.18 |
| | 1259-IMAC | 2.59 | 0.37 |
| | 1259-S1 | 2.25 | 0.45 |
| | 1856-IMAC | 2.37 | 0.42 |
| | 1856-S1 | 2 | 0.51 |
| | 1856-S2 | 1.88 | 0.54 |

**Figure S1. FRET-ratio correlates with FRET efficiencies measured in cell lysates and fluorescence anisotropy reports the assembly density. a, b, c,** FRET efficiency *in vivo* and *in vitro* assessment of control proteins fused to FRET-based proximity biosensor and expressed in *E. coli* (random proteins 665, 1259, 1856; various length glycine-serine linkers; *E. coli* proteins bolA, cspA, iscA, sodA). Controls were analysed by flow cytometry and KDE plots in (**a**) show their FRET ratio distributions — a flow cytometry parameter derived from FRET-channel intensity normalized for donor intensity. Cell lysates of the controls were analysed by time-resolved spectroscopy. Donor lifetime distributions (τ) in (**b**) were extracted with Maximum entropy method (MEM). FRET efficiency ($E_{FRET}$) in (**c**) was calculated from from $E_{FRET}$ ($\tau_{ave}$) = 1 - $\tau_{ave,DA}/\tau_{ave,D}$ where $\tau_{ave}$ is amplitude weighted mean lifetime of the whole $\tau$ distribution of the donor-only (D) and FRET-pair construct (DA). **d, e,** Control proteins 665, 1259 and 1856 were purified (Methods) to determine the FRET efficiency of their high molecular weight oligomeric species. Affinity chromatography elutions (IMAC) and different elution volume fractions (S) from gel chromatography (**f**) were analysed by TCSPC. Lifetime distributions ($\tau$) of purification fractions in (**e**) were used to extract $\tau_{ave}$ and $E_{FRET}$ in (**c**). **f,** Barplot shows steady-state fluorescence anisotropy of the acceptor measured for the random control proteins and the FRET pair connected by glycine-serine linker (GS2). Individual data points represent images with multiple objects. Scatterplot depicts the correlation of the steady-state anisotropy with $E_{FRET}$ calculated from the lifetime of the donor.

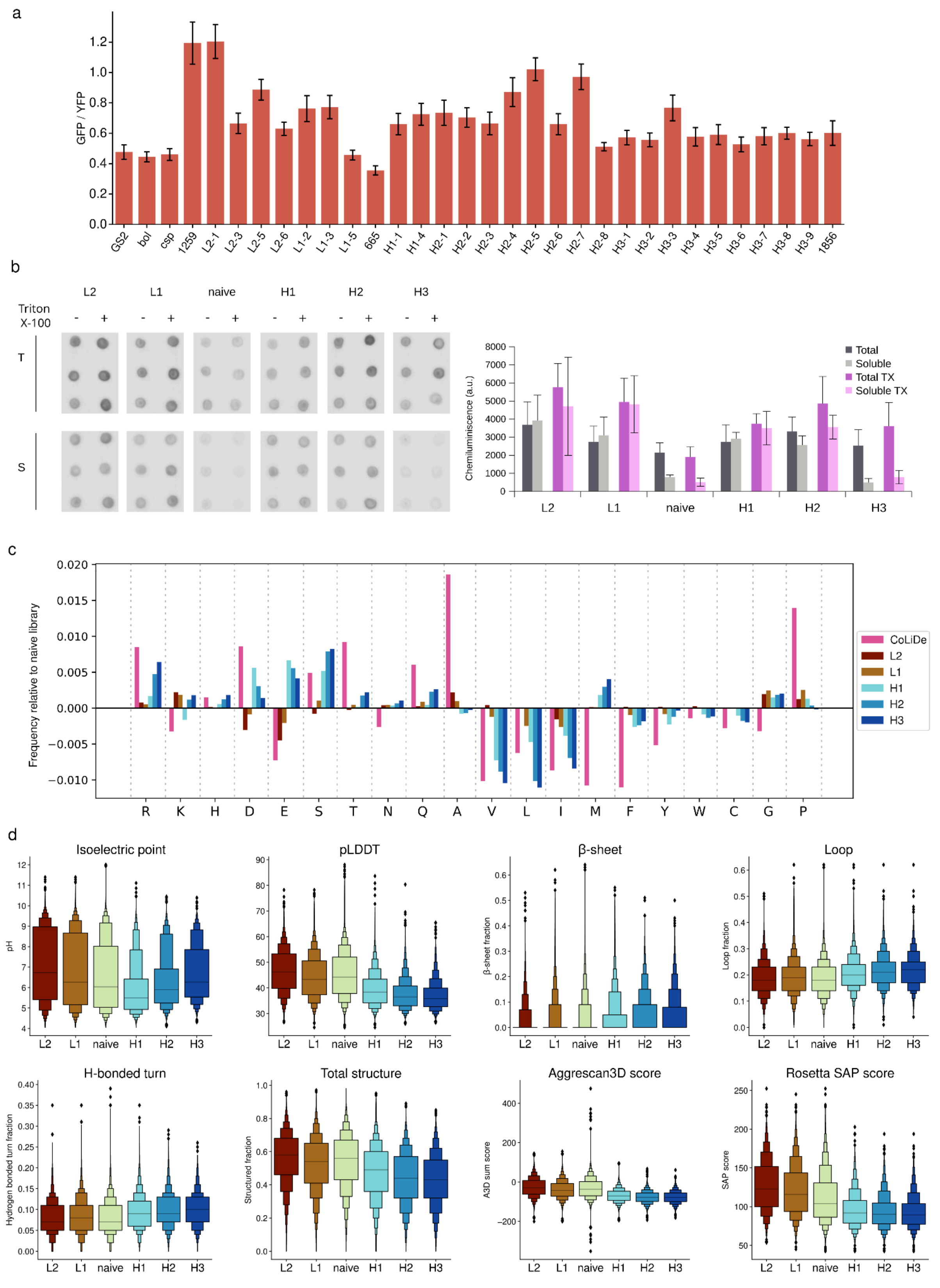


**Figure S2. Chaperone-response assay, experimental solubility and additional properties of sorted subpopulations**

**a,** Response of *ibpA* promoter to the expression of candidate proteins and controls at 37˚C for 1 hour. GFP fluorescence (reporting on the activity of *ibpA*) was normalized to the mVenus fluorescence

(Whiskers show standard deviation from three independent replicates). **b**, Solubility of sorted subpopulations expressed with FLAG-tag fusion in a cell-free system and the effect of Triton X-100. The content of the total and soluble fractions was evaluated by dot-blot assay. **c**, Amino acid composition of the FRET-FACS sorted subpopulations relative to the naive library composition. CoLiDe frequencies were derived from decoding of the degenerate codons used for the 20F109 library design. **d**, Features extracted from sequences (isoelectric point) and features extracted with DSSP4 from structures predicted by AlphaFold2 (□- strand content, total secondary structure, N-C termini distance, turn, bend, loop and SAP score).

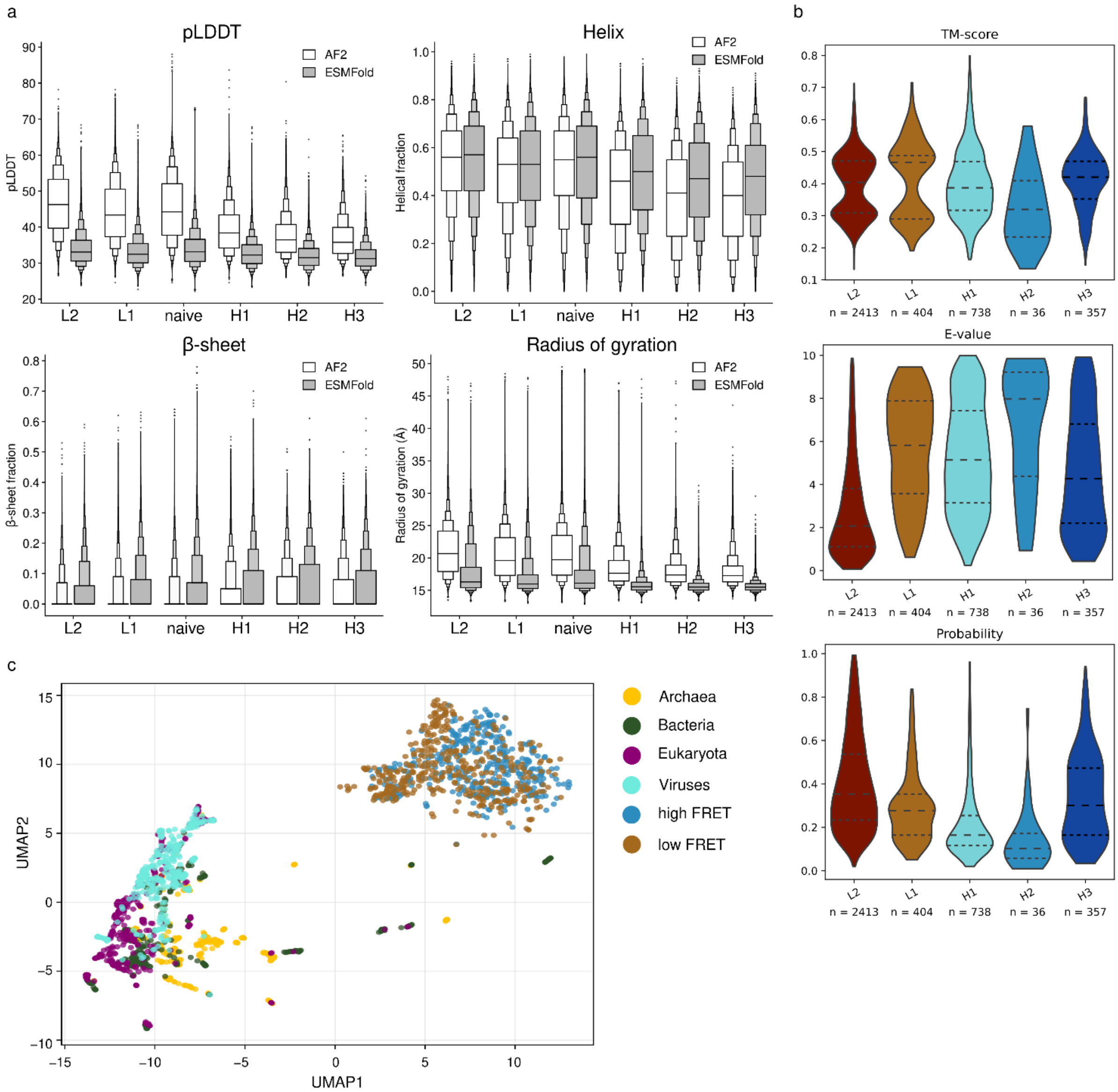


**Figure S3. Comparison of AlphaFold2 and ESMFold predictions, and structural homology search**

**a,** Comparison of AlphaFold2 and ESMFold through selected structure-derived features. **b**, Predicted structures (by ESMFold) of 18 high-abundance candidates from each sorting round were searched against the AlphaFold database using FoldSeek. **c**, Embeddings of length matched natural sequences from different kingdoms with representative sequences from low and high FRET sorting class (per token sequences encoding by ESM-2 model and dimensionality reduction by UMAP).

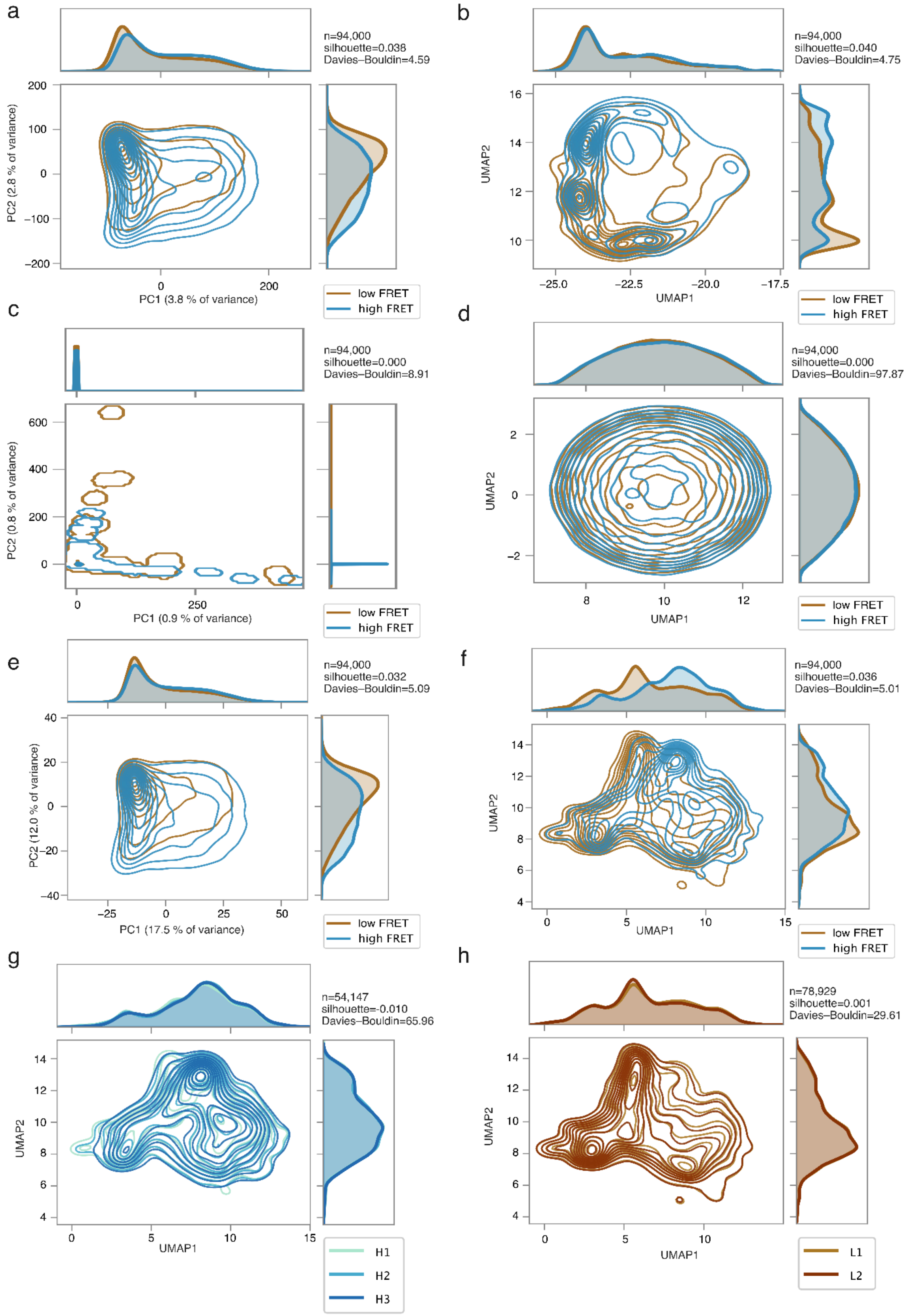


**Figure S4. Different sequence representations plotted into 2D space. a, b,** Contour plots of ESM-2 generated per token sequence representation reduced into 2D space with PCA (**a**) and UMAP projection (**b**). **c, d,** Contour plots of one-hot encoding sequence representations reduced into 2D space with PCA (**c**) and UMAP projection (**d**). **e, f,** Contour plot of ESM-2 generated mean sequence representation reduced into 2D space with PCA (**e**) and UMAP (**f**). **g, h,** Contour plots with sequence representation projection of sorted high-FRET (**g**) and low-FRET (**h**) subpopulations (identical sequence selection as in Figure 2a and Figure S3b). Mean sequence representation generated with ESM-2 and projected with UMAP.

.

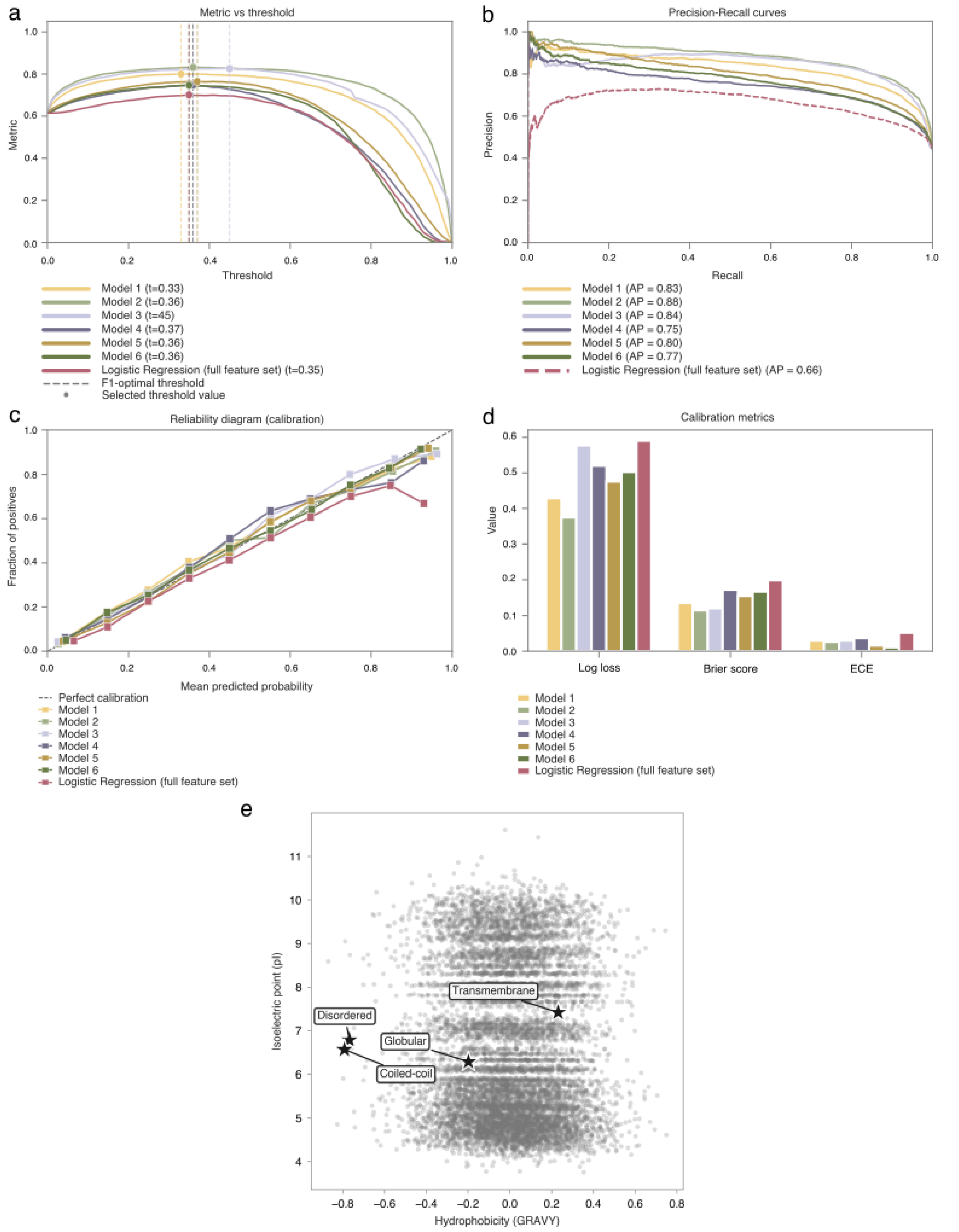


**Figure S5. Model calibration and sequence-composition landscape.** Performance and calibration of six sequence classifiers (Models 1–6) and a logistic regression baseline trained on full composition features ("Logistic Regression (full feature set)"), evaluated on the held-out test set (same stratified split as in Fig. 3). Model colours are consistent across panels. **a,** Metric versus threshold. F1 score as a function of the classification threshold. Solid curves, Models 1–6; dashed red curve, composition logistic regression. Vertical dashed lines mark the F1-optimal threshold for each model (legend: t); filled circles mark the corresponding F1 at that threshold. **b,** Precision–recall curves. Precision versus recall on the test set for

Models 1–6 and the composition logistic regression. Average precision (AP) summarizes overall performance (higher is better). Embedding-based models outperform the composition-only baseline (AP = 0.66), with per-token ESM embeddings performing best (AP = 0.88). All sequence models maintain high precision at low-to-moderate recall, indicating reliable identification of high-FRET variants before errors increase at higher sensitivity. **c,** Reliability diagram. Observed fraction of positives (high-FRET class) versus mean predicted probability in each calibration bin. The diagonal line indicates perfect calibration. Deviation below the diagonal indicates overconfidence; deviation above indicates underconfidence. **d,** Calibration metrics. Grouped bars show Log loss, Brier score and expected calibration error (ECE) for each model (lower is better). Embedding-based models show lower calibration error than the composition-only logistic regression. **e,** Isoelectric point versus hydrophobicity (GRAVY). Grey points, random sample (n = 10,000) from the naive sequence library. Black stars, mean GRAVY and pI for reference sets of disordered (DisProt), coiled-coil (UniProt), globular (SCOPe) and transmembrane (SCOPe) proteins, computed as for the FRET subsets. The high-FRET centroid lies close to the globular reference, whereas the low-FRET centroid falls between the globular/disordered and transmembrane references. Horizontal gaps in the library cloud reflect library-design constraints, not biological absence of sequences in those regions of composition space.
